# Neuronal hypofunction and network dysfunction in a mouse model at an early stage of tauopathy

**DOI:** 10.1101/2024.04.29.591735

**Authors:** Changyi Ji, Xiaofeng Yang, Mohamed Eleish, Yixiang Jiang, Amber Tetlow, Soomin Song, Alejandro Martín-Ávila, Qian Wu, Yanmei Zhou, Wenbiao Gan, Yan Lin, Einar M Sigurdsson

## Abstract

We previously reported altered neuronal Ca^2+^ dynamics in the motor cortex of 12-month-old JNPL3 tauopathy mice during quiet wakefulness or forced running, with a tau antibody treatment significantly restoring the neuronal Ca^2+^ activity profile and decreasing pathological tau in these mice ^1^. Whether neuronal functional deficits occur at an early stage of tauopathy and if tau antibody treatment is effective in younger tauopathy mice needed further investigation. In addition, neuronal network activity and neuronal firing patterns have not been well studied in behaving tauopathy models.

In this study, we first performed in vivo two-photon Ca^2+^ imaging in JNPL3 mice in their early stage of tauopathy at 6 months of age, compared to 12 month old mice and age-matched wild-type controls to evaluate neuronal functional deficits. At the animal level, frequency of neuronal Ca^2+^ transients decreased only in 6 month old tauopathy mice compared to controls, and only when animals were running on a treadmill. The amplitude of neuronal transients decreased in tauopathy mice compared to controls under resting and running conditions in both age groups. Total neuronal activity decreased only in 6 month old tauopathy mice compared to controls under resting and running conditions. Within either tauopathy or wild-type group, only total activity decreased in older wild-type animals. The tauopathy mice at different ages did not differ in neuronal Ca^2+^ transient frequency, amplitude or total activity. In summary, neuronal function did significantly attenuate at an early age in tauopathy mice compared to controls but interestingly did not deteriorate between 6 and 12 months of age.

A more detailed populational analysis of the pattern of Ca^2+^ activity at the neuronal level in the 6 month old cohort confirmed neuronal hypoactivity in layer 2/3 of primary motor cortex, compared to wild-type controls, when animals were either resting or running on a treadmill. Despite reduced activity, neuronal Ca^2+^ profiles exhibited enhanced synchrony and dysregulated responses to running stimulus. Further ex vivo electrophysiological recordings revealed reduction of spontaneous excitatory synaptic transmission onto and in pyramidal neurons and enhanced excitability of inhibitory neurons in motor cortex, which were likely responsible for altered neuronal network activity in this region.

Lastly, tau antibody treatment reduced pathological tau and gliosis partially restored the neuronal Ca^2+^ activity deficits but failed to rescue altered network changes.

Taken together, substantial neuronal and network dysfunction occurred in the early stage of tauopathy that was partially alleviated with acute tau antibody treatment, which highlights the importance of functional assessment when evaluating the therapeutic potential of tau antibodies.

**Highlights:**

- Layer 2/3 motor cortical neurons exhibited hypofunction in awake and behaving mice at the early stage of tauopathy.
- Altered neuronal network activity disrupted local circuitry engagement in tauopathy mice during treadmill running.
- Layer 2/3 motor cortical neurons in tauopathy mice exhibited enhanced neuronal excitability and altered excitatory synaptic transmissions.
- Acute tau antibody treatment reduced pathological tau and gliosis, and partially restored neuronal hypofunction profiles but not network dysfunction.

## Introduction

Accumulation of hyperphosphorylated tau is a major pathological hallmark of Alzheimer’s disease and frontotemporal dementia with tauopathy ^2^. In healthy neurons, tau plays important roles in supporting a variety of neuronal functions, such as axon transport and synaptic signaling ^3^. Under pathological conditions, excessive post-translational modifications of tau lead to its misfolding and aggregation, and thereby compromise normal neuronal function ^4^.

Previous studies have reported abnormal neuronal function in tauopathy mouse models using in-vivo two photon Ca^2+^ imaging ^1,5,6^. Two of them reported a decrease in neuronal activity, as measured by Ca^2+^ transient frequency and total Ca^2+^ activity in two different tauopathy mouse models under anesthesia or quite wakefulness ^5,6^. Our previous study also detected altered neuronal activity in behaving JNPL3 tauopathy mice during resting or treadmill running ^1^. The enhanced neuronal activity during running led to more pronounced abnormal Ca^2+^ profiles compared to resting mice. The mice used in that study were 10-12 month old and had moderate behavior and motor deficits. In JNPL3 mice, tau pathology starts to accumulate at 4-6 months and progresses relatively slowly compared to some other tauopathy mouse models ^7^. Thereby, it allows a wider time window to study tau pathogenesis, the progression of its pathology, and to examine therapeutic efficacy of potential disease-modifying drugs. Since the presence of pathological tau may compromise neuronal function much earlier than the onset of behavior deficits, we sought to identify early neuronal deficits in young JNPL3 mice at the onset of tau pathology at 6 months of age, compared to mice at 12 months of age.

Different neuron types exhibit distinct patterns of activity during motor behaviors. For example, some neurons are activated while others are suppressed during running ^8,9^. This segregation of neuronal firing patterns underlies circuitry mechanisms that are critical for information processing, movement initiation and execution ^8–10^. Neuronal network function can involve rhythmic and synchronized firing ^11^, and coherent patterns are essential for cognitive function ^11^. However, excessive synchronization may be pathological, as seen in seizures associated with epilepsy and Alzheimer’s disease ^12,13^. It remains unclear whether and how tau pathology impacts neuronal network activity. During motor behaviors, excitatory pyramidal neurons in the primary motor cortex increase their activity, and exhibit sequential activation that is stabilized after motor learning ^8–10^. Inhibitory neurons display more diverse responses ^8,14^, and inhibition shapes the activity profiles of pyramidal neurons by tuning their responsiveness to motor stimuli ^8,10,14^. Whether the neuronal firing pattern during motor behaviors changes under pathological conditions like tauopathy warrants further investigation.

In this study, we performed in vivo two-photon Ca^2+^ imaging in 6 and 12 month old JNPL3 mice and their age-matched wild-type (WT) controls when mice were either awake and resting, or performing a running task on the treadmill (Figure 1A-B). In 6 month old mice, we observed significant decrease of neuronal activity in JNPL3 mice compared to WT controls, as indicated by the frequency of Ca^2+^ transients at the running condition, as well as by Ca^2+^ transient amplitude and integrated total Ca^2+^ activity at both resting and running conditions. In 12 month old cohorts, only Ca^2+^ transient amplitude decreased significantly in JNPL3 mice compared to age-matched WT controls at both resting and running conditions. In addition, age-dependent decrease of total Ca^2+^ activity was observed in WT mice, but not JNPL3 mice. Furthermore, we examined Ca^2+^ activity profiles and neuronal population activity in the younger cohort, which was not explored in our previous study ^1^. The proportion of hypoactive neurons was significantly larger in JNPL3 mice, and running related responses were largely suppressed. This reduction in neuronal Ca^2+^ activity was likely associated with decreased excitatory synaptic connections. Nevertheless, motor cortical neurons in JNPL3 mice exhibited higher synchronization compared to those in WT mice. This change might be related to enhanced inhibitory neuronal activity. Lastly, acute treatment of the tauopathy mice with a tau monoclonal antibody partially restored the decreased neuronal activity but not altered network activity, and the antibody decreased soluble phosphorylated tau, increased insoluble total tau and attenuated gliosis in the mice.

**Figure 1.**
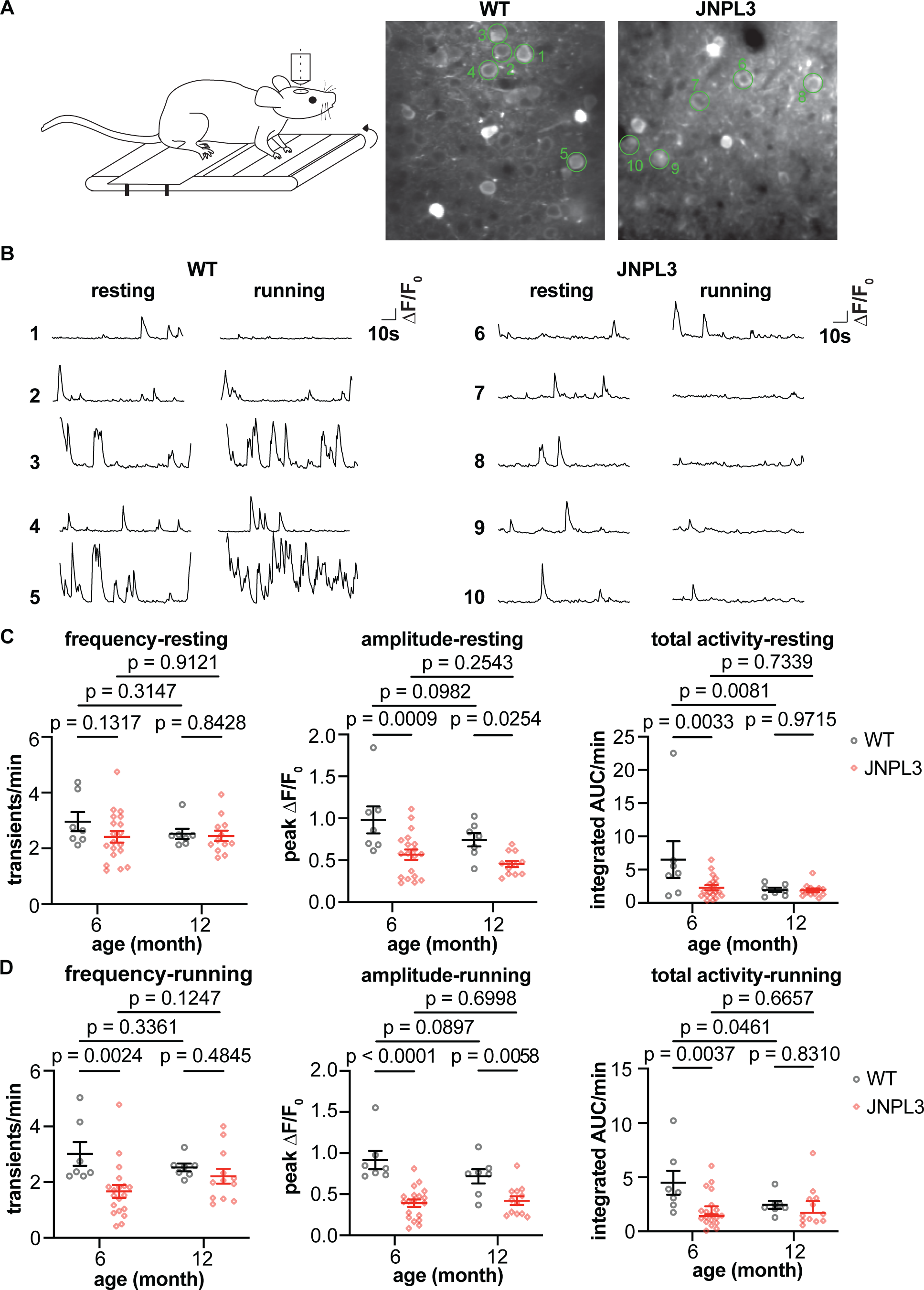
Decreased somatic Ca^2+^ activity in L2/3 motor cortex in 6 month old JNPL3 mice at early stage of tauopathy. **(A)** Schematic representation of two-photon imaging in head-fixed, awake and behaving mice. GCaMP6s was expressed in motor cortex layer 2/3 neurons. Representative images of AAV-syn-GcaMP6s expression are shown in the right panel with green circles outlining analyzed neurons. **(B)** Representative Ca^2+^ transient traces in motor cortex layer 2/3 neurons when mice were either resting or running. F_0_: fluorescence intensity at baseline, ΔF: fluorescence intensity per time point minus fluorescence intensity at baseline. **(C)** Ca^2+^ activity frequency (transients/min), Ca^2+^ amplitude (peak ΔF/F_0_), and total Ca^2+^ activity (integrated area under the curve (AUC)/minute) were analyzed in 6 month or 12 month old mice when animals were resting on the treadmill. **(D)** The same parameters were analyzed in the same 6 and 12 month old mice when animals were forced to run on the treadmill. The 6 month old group consisted of 7 WT and 19 JNPL3 mice, and the 12 month group consisted of 7 WT and 12 JNPL3 mice. Each data point represents one animal. Two-way ANOVA with Fisher’s LSD multiple comparisons test in (C) and (D).

## Results

### Altered somatic Ca^2+^ activity in JNPL3 mice during early vs later stage of tauopathy

In this study, we first determined if Ca^2+^ abnormalities can be detected before the development of robust tau pathology. We followed the similar experimental paradigm as described in our previous study ^1^ and performed in vivo two-photon Ca^2+^ imaging in L2/3 motor cortical region when animals were resting or forced to run on a treadmill. We analyzed neuronal Ca^2+^ profiles in both 6 month old and 12 month old JNPL3 mice, and compared to their age-matched WT controls.

In resting 6 month old mice, the amplitude of Ca^2+^ transients and total Ca^2+^ activity significantly decreased in JNPL3 mice compared to age-matched wild-type (WT) controls. However, the frequency of somatic Ca^2+^ transients did not change (Figure 1C). When these mice were running on the treadmill, all these three parameters significantly decreased in JNPL3 mice compared to WT controls (Figure 1D). In 12 month old JNPL3 mice, only the amplitude of Ca^2+^ transients decreased under both resting and running conditions, whereas the frequency of Ca^2+^ transients and total Ca^2+^ activity did not differ significantly between JNPL3 and age-matched WT mice (Figure 1C and 1D). Interestingly, the total Ca^2+^ activity decreased significantly in 12 month old WT mice compared to 6 month old WT mice. However, no significant difference was observed in other Ca^2+^ activity parameters between these two age groups within either WT or JNPL3 groups. Overall, awake and behaving JNPL3 mice at an early stage of tauopathy exhibited a pronounced decrease of neuronal activity in motor cortex compared to age-matched WT mice. This attenuation of neuronal activity did not further deteriorate from 6 to 12 months of age. This difference may in part relate to a lower neuronal activity in the older mice, regardless of genotype.

### Neuronal hypofunction in 6 month old JNPL3 mice

To gain further insights into abnormal neuronal activity observed in 6 month old JNPL3 mice, we further looked into the pattern of Ca^2+^ activity at the neuronal level, which had not been thoroughly examined in our previous study ^1^. There was a significant change of distribution of Ca^2+^ transient frequency in JNPL3 mice compared to WT mice, during both resting and running conditions (Figure 2A). We classified these neurons into three categories based on their Ca^2+^ activity frequency. Neurons exhibiting less than two transients per minute were considered to be hypoactive, whereas those with more than six transients per minute were categorized as hyperactive ^6,18–20^. Neurons with two to six transients per minute were marked to have normal activity.

**Figure 2.**
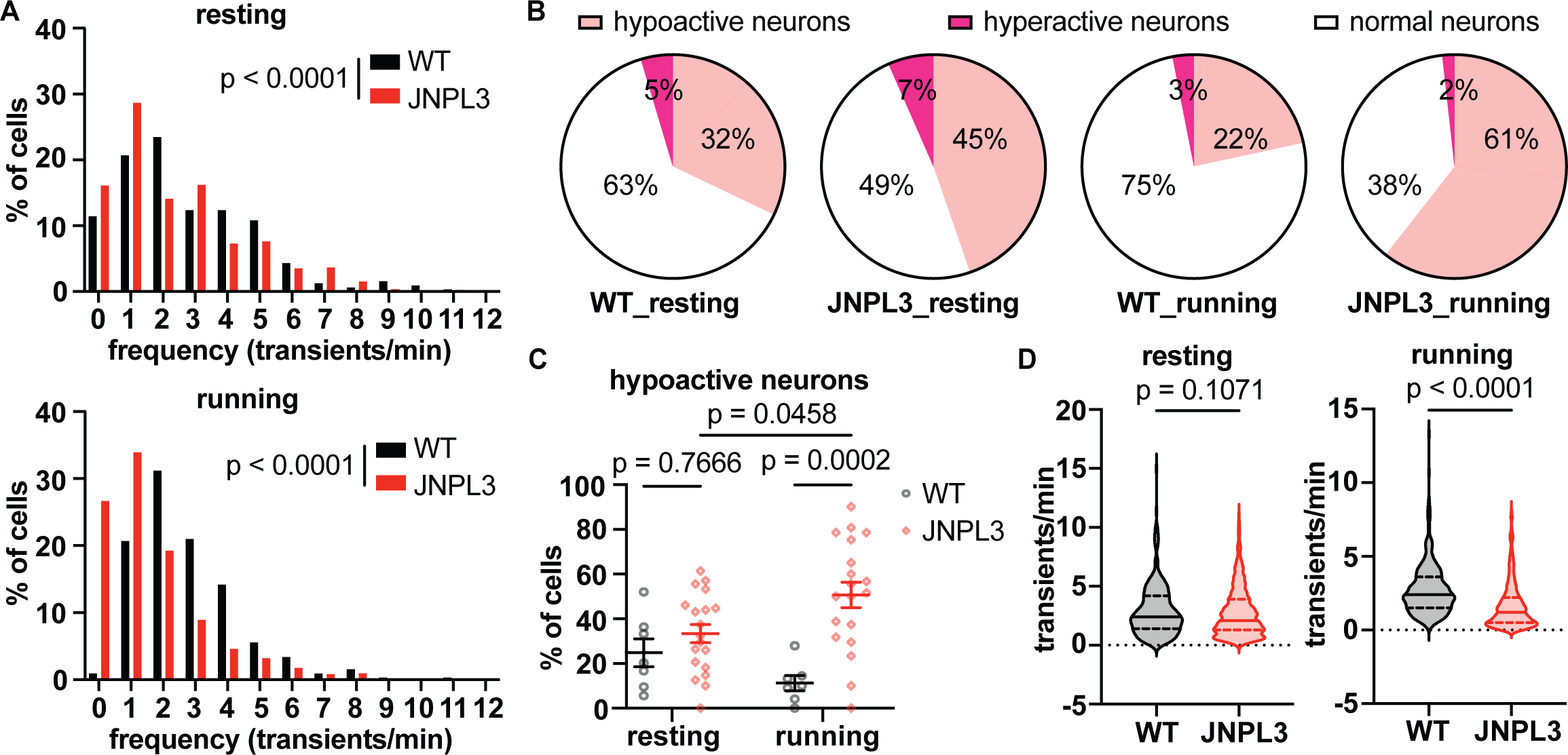
Increased hypoactive neurons in running JNPL3 L2/3 motor cortex. **(A)** Further analysis of all neurons showed altered frequency distribution of Ca^2+^ transients in JNPL3 mice when animals were either resting or running on the treadmill, compared to WT mice. **(B)** The fraction of hypoactive neurons increased in both resting and running JNPL3 mice, compared to WT mice. Hypoactive neurons: <2 transients/min; hyperactive neurons: >6 transients/min; normal neurons: 2-6 transients/min. A total of 325 neurons from 7 WT animals and 852 neurons from 19 JNPL3 mice were analyzed. **(C)** The fraction of hypoactive neurons in each animal (represented by a data point). **(D)** Ca^2+^ transient frequency in active neurons (>0 transients/min). While resting, a total of 311 neurons from WT animals and 737 neurons from JNPL3 mice were analyzed. While running, a total of 325 neurons from WT animals and 800 neurons from JNPL3 mice were analyzed. Two-way ANOVA with Tukey’s multiple comparisons test in (C). Mann-Whitney U test in (D).

In resting WT mice, 32% of neurons were hypoactive, 63% were normal, and 5% were hyperactive (Figure 2B). When these mice were running, 22% of neurons were hypoactive, 75% had normal activity, and 3% were hyperactive (Figure 2B). In age-matched JNPL3 mice, the fraction of hypoactive neurons increased to 45%, and hyperactive neurons increased to 7% when animals were at rest (Figure 2B). As a result, the fraction of neurons with normal activity decreased to 49% (Figure 2B). When JNPL3 mice were running on the treadmill, the fraction of hypoactive neurons increased further to 61%, hyperactive neurons decreased to 2%, and normal neurons went down to 38% (Figure 2B).

We further analyzed the fraction of hypoactive neurons in each animal to confirm that this result was not confounded by dominant changes in certain animals. Consistently, running JNPL3 mice had a larger fraction of hypoactive neurons than running WT controls (JNPL3 vs WT: 51% vs 11%). Moreover, the fraction of hypoactive neurons in JNPL3 mice during running was also larger than the fraction when these animals were at resting state (running vs resting: 51% vs 33%) (Figure 2C). With regard to the average frequency of Ca^2+^ transients in active neurons, it decreased in running but not in resting JNPL3 mice, compared to WT controls (Figure 2D).

Consistent with our previous study in 12 month old animals ^1^, the peak amplitude of Ca^2+^ transients and total Ca^2+^ activity in active neurons decreased in 6 month old JNPL3 mice compared to WT controls, when animals were either resting or running on the treadmill (Figure S1A, S1C, S1E, and S1G). There was a left shift of cumulative frequency distribution of peak Ca^2+^ amplitude and total Ca^2+^ activity (Figure S1B, S1D, S1F, and S1H). Taken together, JNPL3 mice had lower neuronal activity compared to their age-matched WT controls.

### Altered neuronal network activity in 6 month old JNPL3 mice

Given the decreased neuronal activity in the JNPL3 mice, we further assessed the impact of tau pathology on neuronal circuitry in the primary motor cortex. We analyzed the correlations of neuronal Ca^2+^ dynamics, which reflect the strength of neuronal interactions within a given population ^5,21^. First, we calculated Pearson’s correlation coefficients between a given neuron and all other neurons in the field of view, which resulted in a correlation matrix (Figure 3A and 3D). Compared to WT mice, Ca^2+^ activity correlations were more prominent in JNPL3 mice during both resting and running periods (Figure 3A and 3D). Subsequently, we calculated a correlation index for a given neuron by averaging the Pearson’s correlation coefficients of that neuron paired with all other neurons in the field of view. The correlation index significantly increased at the neuronal (Figure 3B) and animal (Figure 3C) level in resting JNPL3 mice. A significant increase of the correlation index was also observed in running JNPL3 mice at the neuronal level (Figure 3E), but not at the animal level (Figure 3F). These results suggested increased synchrony of neuronal activity in JNPL3 mice, especially at the resting period.

**Figure 3.**
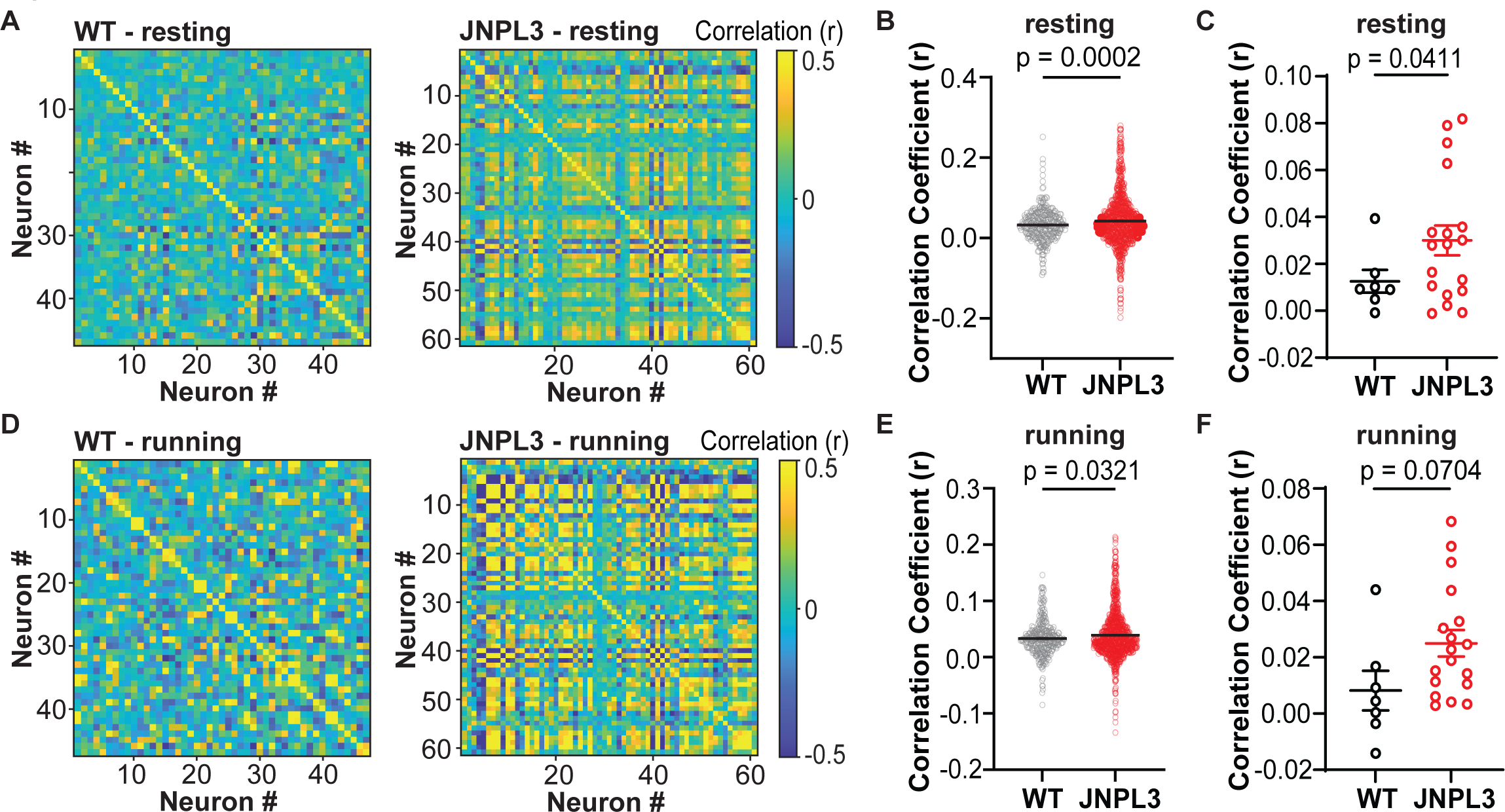
Increased neuronal activity synchrony of L2/3 motor cortex in JNPL3 mice. **(A)** Correlation coefficient matrix of Ca^2+^ activity in a WT or JNPL3 mouse when animals were resting. In a given field of view (FOV), Ca^2+^ activity from every two neurons were compared in pairs, and accessed by Pearson’s correlation coefficiency. **(B)** Ca^2+^ activity correlation in all neurons in resting animals. A correlation index was calculated for each neuron by averaging the Pearson’s correlation coefficient of a given neuron with all other neurons in the same FOV. Each dot represented a neuron. Significant increase of Ca^2+^ activity correlation was observed in resting JNPL3 mice compared to WT mice. **(C)** Ca^2+^ activity correlation in resting WT and JNPL3 mice. Each dot represented the mean correlation index of all neurons in a given animal. **(D)** Same analyses as (A) in running animal. **(E)** Same analysis as (B) in running animals. Ca^2+^ activity correlation significantly increased in running JNPL3 mice compared to WT controls. **(F)** Same analysis as (C) in running animals. Pearson’s correlations in (A) and (D). Unpaired t test in (B), (C), (E), and (F).

Different neurons respond to the same stimuli in different yet organized ways, which may reflect circuit mechanisms during information processing ^8,10^. We next analyzed the responses of neuronal Ca^2+^ activity during running relative to resting (Figure 4A and 4B). Neurons were categorized into four types based on their responses to running relative to resting: 1) activated activity; 2) suppressed activity; 3) mixed response (both activated and suppressed activity), or 4) no change (Figure 4C). In WT mice, 37% of neurons were activated, 15% were suppressed, 41% had mixed responses, and 7% did not respond to running. In JNPL3 mice, 23% of neurons were activated, 46% showed suppressed activity, 21% had mixed responses, and 10% did not respond to running. In running JNPL3 mice compared to WT mice, the fractions of activated neurons decreased significantly, whereas the fractions of suppressed neurons increased significantly (Figure 4D and 4E). Although not statistically significant, there was a trend towards a decrease in the fraction of mixed responsive neurons in JNPL3 mice compared to WT mice (Figure 4E). Additionally, the fraction of neurons that lacked response to running remained the same between JNPL3 and WT mice (Figure 4E). Overall, neuronal activity was suppressed in running JNPL3 animals, which further supports the hypofunction of motor cortical neurons in JNPL3 tauopathy mice at 6 months of age. Given the diverse responses of neuronal activity to running, Ca^2+^ activity correlation was assessed separately in neurons with either activated or suppressed activity when animals were running. In neurons that were activated during running, the Ca^2+^ activity correlation increased significantly in JNPL3 mice compared to WT mice (Figure 5A and 5B). In contrast, the correlation coefficiency decreased significantly in neurons with suppressed Ca^2+^ activity when JNPL3 mice performed running tasks compared to WT mice (Figure 5C and 5D). Taken together, these data suggested that pathological tau altered the engagement pattern of local neuronal circuitry in primary motor cortical L2/3 region of JNPL3 mice during running behaviors.

**Figure 4.**
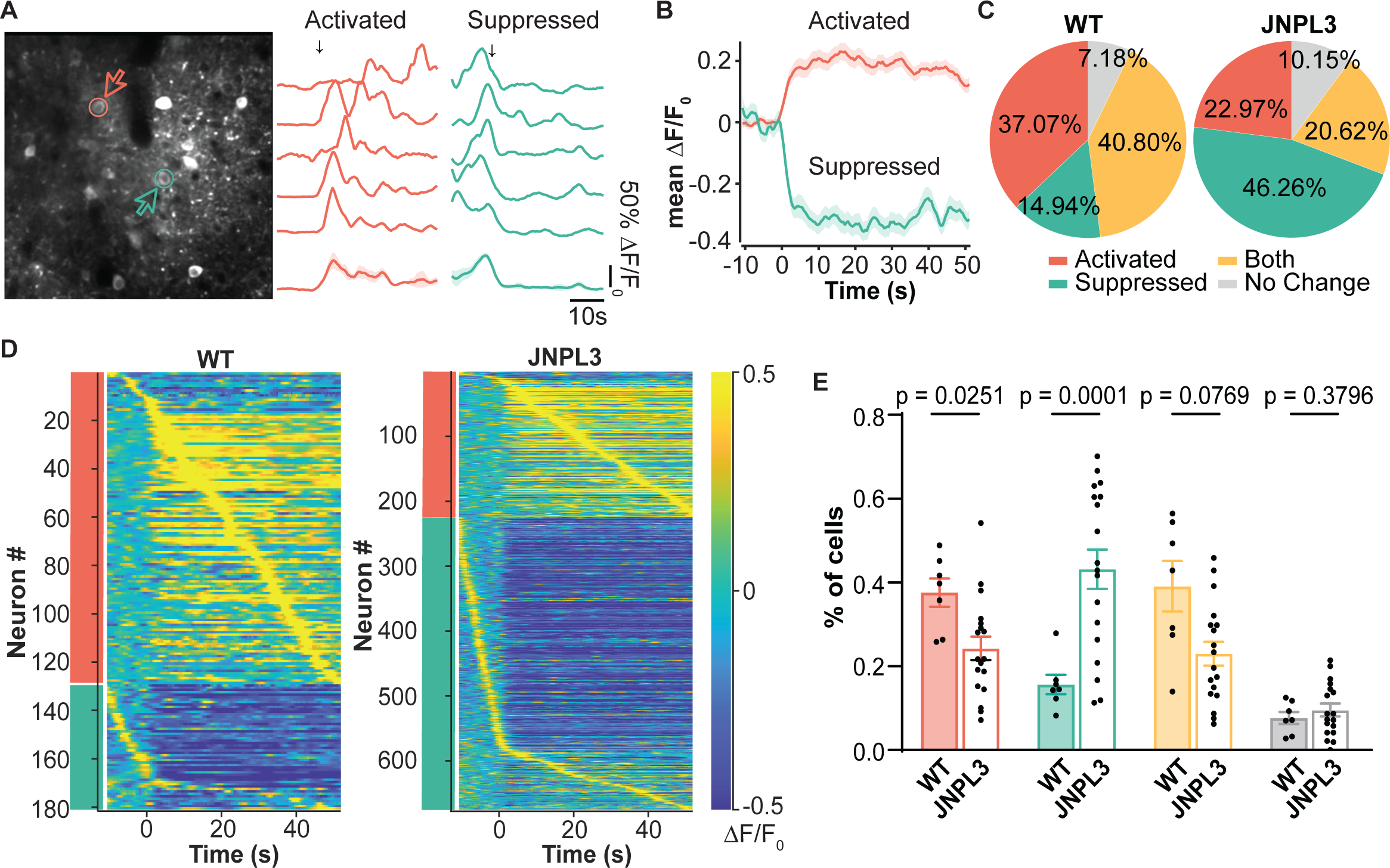
Altered neuronal responses to running of L2/3 motor cortical neurons in JNPL3 mice. **(A)** Representative Ca^2+^ transient traces from two neurons of a WT mouse showed either activated (red) or suppressed (green) activity in response to running. Ca^2+^ traces (normalized ΔF/F) from five trials were aligned to the onset of treadmill rotation as indicated by the arrow. The average trace of all five trials is shown at the bottom. The shade represents SEM. Vertical scale bar equals 50% maximum normalized ΔF/F_0_, horizontal scale bar represents 10 s. **(B)** Average traces of all neurons exhibiting activated or suppressed responses to running in a WT mice. Solid line represents mean value, and shaded region shows SEM. **(C)** Neurons were categorized into four types according to their responses to running. In WT mice (348 neurons, 7 animals), 37.07% neurons were activated (red), 14.94% were suppressed (green), 40.8% showed both activated and suppressed responses (yellow), and 7.18% had no change (grey). In JNPL3 mice (975 neurons, 18 animals), 22.97% neurons were activated, 46.26% were suppressed, 20.62% shows both activated and suppressed responses, and 10.15% had no change. **(D)** Activity pattern of neurons with activated (red bar) or suppressed (green bar) response to running in WT (left) and JNPL3 (right) mice. The average activity of each neuron from five trials was aligned to the onset of treadmill, and then sorted by the time taken to their peak activity. (**E)** The fraction of each neuronal responses type in each animal when mice were running. The ratio of activated neurons decreased while those with suppressed activity increased in JNPL3 mice compared to WT mice. Each dot represented an animal. Multiple unpaired t test.

**Figure 5.**
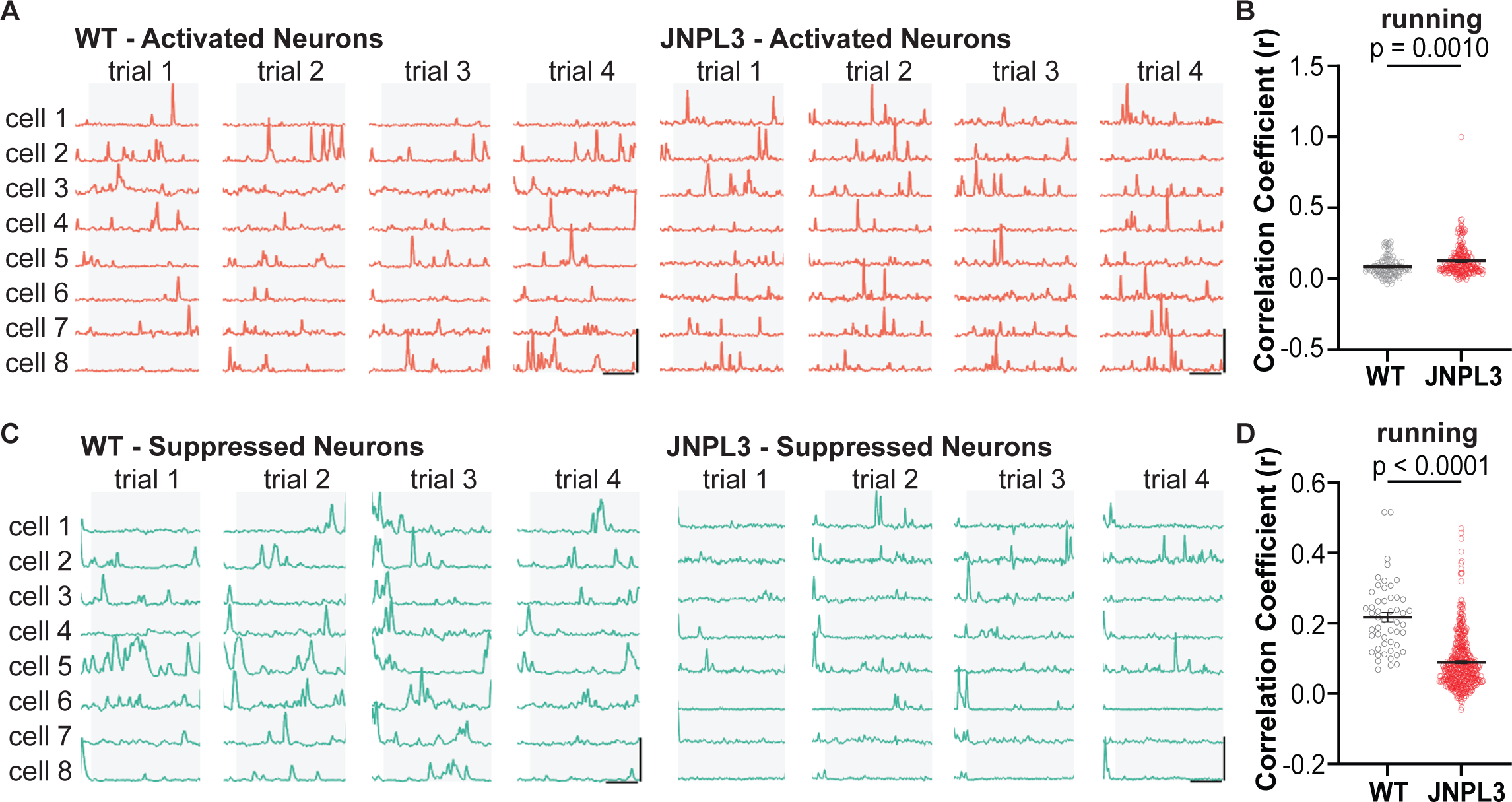
Differential changes of neuronal activity correlation in neurons with activated or suppressed response to running in JNPL3 mice. **(A)** Representative Ca^2+^ trace of neurons with activated responses in a WT and a JNPL3 mouse in running trials. Grey background/box depicted running periods whereas clear background/box denoted resting periods. **(B)** Ca^2+^ activity correlation in neurons with activated response to running. Same analyses of correlation index were performed as (Figure 3B). A total of 129 neurons (7 animals) from WT and 224 neurons from JNPL3 (18 animals) were included. **(C)** Representative Ca^2+^ trace of neurons with suppressed responses during running in a WT and a JNPL3 mouse. **(D)** Ca^2+^ activity correlation in neurons with suppressed response to running. Same analyses of correlation index were performed as (Figure 3B). A total of 52 neurons (7 animals) from WT and 451 neurons from JNPL3 (18 animals) were included. Unpaired t test in (H), (J).

### Increased neuronal excitability and changed synaptic transmissions in 6-month-old JNPL3 mice

To further dissect the mechanisms underlying hypofunction of motor cortical neurons and disrupted network activity in JNPL3 mice, we assessed the intrinsic firing ability and connectivity of L2/3 motor cortical neurons through ex vivo patch-clamp experiments. Neurons exhibiting fast-spiking characteristics (FS), which most likely were parvalbumin expressing interneurons, showed enhanced excitability in JNPL3 brain slices (Figure 6A and 6B). These neurons fired action potentials with larger afterhyperpolarization potential (AHP) compared to WT controls (Figure 6C and 6D). Other parameters related to intrinsic firing properties, such as input resistance, cellular capacitance and rheobase current, were comparable between JNPL3 and WT mice (Table 2). Interestingly, neurons without fast-spiking properties (non-FS) demonstrated significantly increased excitability in JNPL3 mice compared to those in WT mice (Figure 6E and 6F). Similar to FS neurons, the amplitude of AHP in non-FS neurons also increased in JNPL3 mice (Figure 6G and 6H). The resting membrane potential was depolarized in non-FS neurons of JNPL3 mice (WT, -75.07±1.37 mV; JNPL3 mice, -69.07±2.19 mV), which fired action potentials at lower rheobase currents (WT = 283.85 ± 24.74 pA; JNPL3 mice = 113.33 ± 18.91 pA). Consistently, non-FS neurons in JNPL3 mice had decreased cellular capacitance and increased input resistance (Table 2). These non-FS neurons were very likely pyramidal excitatory neurons given their abundance in L2/3 motor cortex, although a minor fraction of them may be somatostatin or vasoactive intestinal peptide expressing inhibitory neurons. Overall, L2/3 motor cortical neurons exhibited enhanced intrinsic firing ability in JNPL3 mice compared to WT mice.

**Figure 6.**
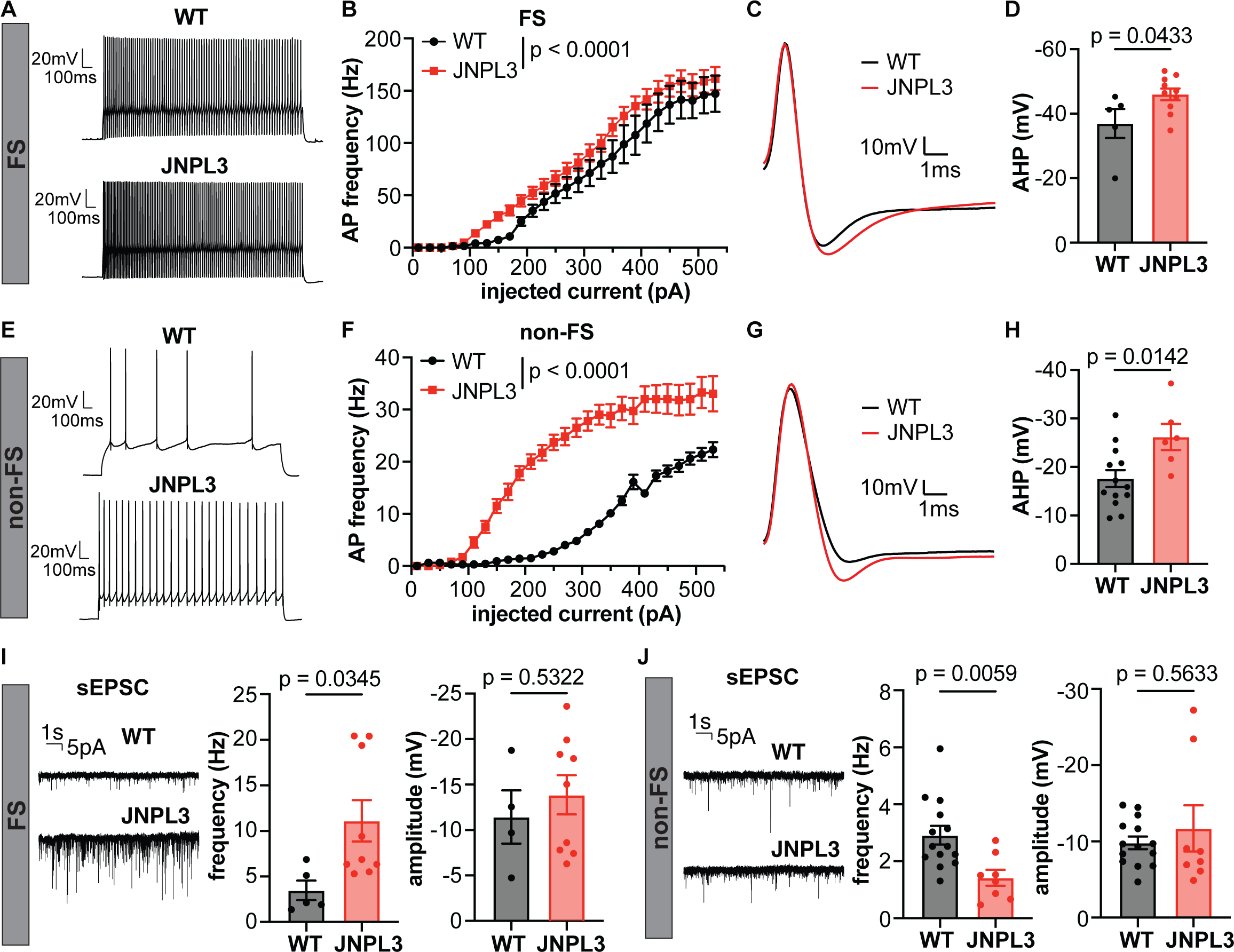
Increased excitability and altered excitatory synaptic transmission of L2/3 motor cortical neurons in JNPL3 mice. **(A)** Representative action potential (AP) firing pattern in a fast-spiking (FS) neurons from a WT or JNPL3 mouse. Neurons were injected with a 290pA depolarization current for 1s. **(B)** Frequency current (F/I) curve of FS neurons in WT or JNPL3 brain slices. Slightly, but significantly left shift of F/I curve in JNPL3 FS neurons compared to WT controls. **(C)** Representative AP waveforms fired by FS neurons with minimal current injection. **(D)** Afterhyperpolarization (AHP) potential significantly increased in JNPL3 FS neuron. **(E)** Exemplar traces of AP firing pattern in non-fast-spiking (non-FS) neurons injected with a 290pA depolarization current for 1s. **(F)** F/I curve of non-FS neurons in WT or JNPL3 brain slices. Significantly left-shift of F/I curve in JNPL3 non-FS neurons compared to WT controls. **(G)** Representative AP waveforms fired by non-FS neurons with minimal current injection. **(H)** Afterhyperpolarization (AHP) potential significantly increased in JNPL3 non-FS neurons. **(I)** Frequency and amplitude of spontaneous excitatory post-synaptic current (sEPSC) in FS neurons. Significant increase of sEPSC frequency but not amplitude was observed in JNPL3 FS neurons. **(J)** Frequency and amplitude of sEPSC in non-FS neurons. Significant decrease of sEPSC frequency but not amplitude was observed in JNPL3 non-FS neurons. Two-way ANOVA followed by Sidak’s multiple comparisons test in (B) and (F). Unpaired t test in (D), (H), (I), and (J).

**Table 1.** Statistical methods used in figures.

| Figure Name |  | Statistical Methods |
| --- | --- | --- |
| Figure 1 | C, D | Two-way ANOVA with Fisher's LSD multiple comparisons test |
| Figure 2 | A | Kolmogorov-Smirnov test |
|  | C | Two-way ANOVA with Tukey's multiple comparisons test |
|  | D | Mann-Whitney U test |
| Figure 3 | A, D | Pearson's correlations |
|  | B, C, E, F | Unpaired t test |
| Figure 4 | E | Multiple unpaired t test |
| Figure 5 | B, D | Unpaired t test |
| Figure 6 | B, F | Two-way ANOVA with Sidak's multiple comparisons test |
|  | D, H, I, J | Unpaired t test |
| Figure 7 | B, C, D, F | Unpaired t test |
| Figure 8 | A - F | Wilcoxon matched-pair signed rank test |
| Supplemental figure 1 | A, C, E, G | Mann-Whitney U test |
|  | B, D, F, H | Kolmogorov-Smirnov test |
| Supplemental figure 2 | B, C, E, F, H, I | Unpaired t test |
| Supplemental figure 3 | B | Kolmogorov-Smirnov test |
|  | D | Multiple unpaired t test |
|  | F | Unpaired t test |
| Supplemental figure 4 | C | Multiple unpaired t test |
|  | E | Unpaired t test |
| Supplemental figure 5 | A | Wilcoxon matched-pair signed rank test |

**Table 2.** Electrophysiological parameters of motor cortical L2/3 neurons in WT and JNPL3 mice. FS, fast-spiking neurons; non-FS, non fast-spiking neurons; Rm, input resistance; Cm, cell capacitance; Rheobase current, the minimal inject current to elicit an action potential.

| Cell type | Genotype | Resting membrane potential (mV) | Rm (MΩ) | Cm (pF) | Rheobase current (pA) |
| --- | --- | --- | --- | --- | --- |
| FS | WT (n = 5) | -73.05 ± 0.71 | 153.80 ± 26.98 | 28.8 ± 2.87 | 159.80 ± 51.75 |
| | JNPL3 (n = 10) | -70.4 ± 1.15 | 191.30 ± 33.22 | 26.9 ± 2.63 | 160.90 ± 32.57 |
|  | p-value (unpaired t test) | 0.1496 | 0.4774 | 0.6632 | 0.9861 |
| non-FS | WT (n = 14) | -75.07 ± 1.37 | 78.36 ± 17.69 | 86.4 ± 8.54 | 283.85 ± 24.74 |
| | JNPL3 (n = 6) | -69.07 ± 2.19 | 151.00 ± 14.20 | 46.8 ± 5.95 | 113.33 ± 18.91 |
|  | p-value (unpaired t test) | 0.0291 | 0.0216 | 0.0101 | <0.0001 |

Furthermore, we assessed synaptic transmissions in FS and non-FS neurons by recording postsynaptic currents. The frequency of spontaneous excitatory post-synaptic current (sEPSC) in FS neurons significantly increased in JNPL3 compared to WT mice (Figure 6I), suggesting enhanced excitatory synaptic transmission in these FS neurons in the presence of mutant tau. In contrast, the frequency of sEPSC in non-FS neurons significantly decreased in JNPL3 mice compared to WT mice (Figure 6J), suggesting reduced excitatory transmission in these cells. The amplitude of sEPSC in either FS or non-FS neurons remained similar between JNPL3 and WT mice.

Taken together, these data indicated impaired excitatory synaptic connections in non-FS neurons in the L2/3 motor cortex of JNPL3 mice compared to WT mice, which may lead to homeostatic upregulation of their excitability ^22^. In contrast, FS neurons received enhanced excitatory synaptic transmission, which likely resulted from increased excitability of pyramidal neurons, and further led to their own enhanced firing ability.

### Reduced pathological tau and gliosis in 6 month old JNPL3 mice after acute tau antibody 8B2 treatment

Previous study from our group showed that a tau monoclonal antibody 4E6 that targets phosphorylated tau-Ser396/Ser404 (pS396/S404) partially restored abnormal Ca^2+^ activity and decreases pathological tau in 10-12 month old JNPL3 mice ^1^. The improvement in Ca^2+^ activity correlated with decreased soluble pathological tau ^1^. Another tau mAb, 8B2, also targets pS396/S404, but has a different binding profile than 4E6 ^17^. In the next step, we examined whether 8B2 also reduced pathological tau levels and rescued abnormal Ca^2+^ activity in motor cortical neurons in awake and behaving JNPL3 mice. JNPL3 mice were imaged at Day 0, followed by intravenous injections of two doses of either 8B2 (100 μg per dose) or IgG1 control 3 days apart (Day 1 and 4), followed by imaging again at Day 7 and brain extraction at Day 8 (Figure 7A). Ca^2+^ activity profiles were assessed before and after the acute antibody treatment, and brain tau levels at the end of the study. A subset of mice was injected intravenously with 8B2 or control IgG1 labeled with VivoTag 680XL to assess 8B2 target engagement. First, we examined target engagement of 8B2 by immunostaining brain sections collected after imaging sessions. Substantially more 8B2 was detected in L2/3 motor cortical neurons compared to IgG1 control (Figure 7B). In addition, immunostaining results revealed that PHF1 and labeled-8B2 overlapping signals were significantly more than PHF1 and labeled IgG1 signals (Figure 7C). Similarly, CP27 and labeled 8B2 signals were also significantly more than CP27 and labeled IgG1 signals (Figure 7D). Furthermore, the 8B2 treatment significantly decreased soluble phosphorylated tau Ser396/Ser404 (pS396/S404, PHF1) and phosphorylated tau Thr231 (pT231) epitopes, but did not affect total tau levels (CP27) in the low-speed supernatant (LSS) of brain homogenates (Figure 7E and 7F). We further applied sarkosyl extraction to enrich insoluble tau aggregates from LSS. Treatment with 8B2 significantly increased sarkysol-insoluble (SP) total tau detected by CP27 (Figure 7E and 7F). Overall, acute 8B2 treatment decreased soluble phosphorylated tau and increased insoluble total tau in JNPL3 mice motor cortex.

**Figure 7.**
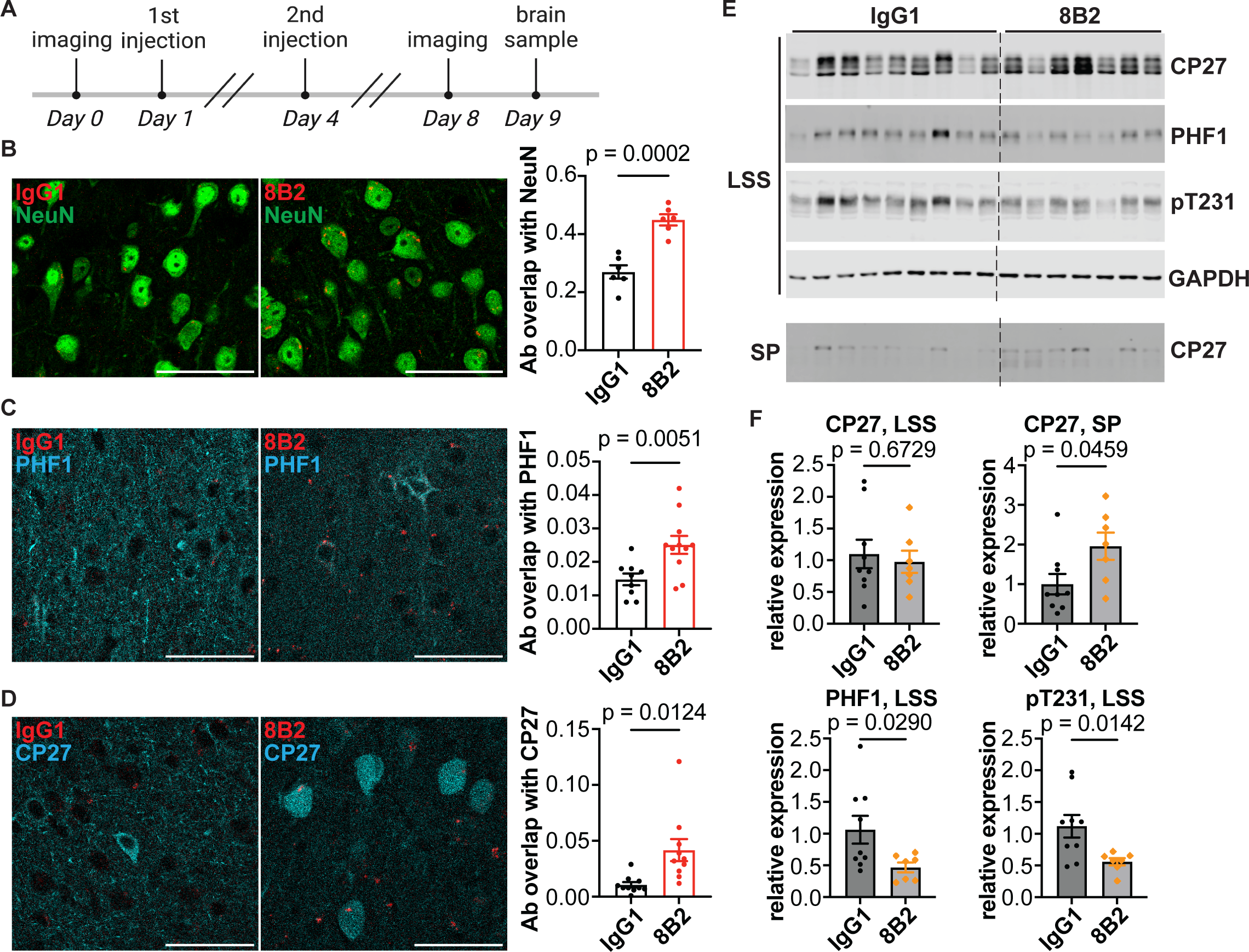
Tau antibody 8B2 is uptake into neurons and colocalized with tau and phosphorylated tau in the motor cortex of 6-month-old JNPL3 mice. **(A)** Schematic illustration of the imaging and antibody injection protocol. **(B)** Tau antibody 8B2 or control IgG1 labeled with VivoTag 680XL (red) were injected intravenously and the brains were extracted for analyses. Brain sections were stained with NeuN antibody. NeuN and 8B2 colocalization was more prominent than NeuN and IgG1 colocalization. **(C)** Brain sections stained with PHF1 antibody. **(D)** Brain sections stained with CP27 antibody. More 8B2 was colocalized with PHF1 or CP27 compared to control IgG1. **(E)** Low speed soluble (LSS) protein fraction from brains lysates of the same JNPL3 mice as in (B) were blotted with CP27, PHF1 and pT231 and GAPDH antibodies. Insoluble protein (SP) fraction of LSS was blotted with CP27. CP27 recognized total human tau. PHF1 and pT231 recognized phosphorylated tau S396/S404 and T231, respectively. GAPDH was a loading control. **(F)** Quantifications of tau protein. LSS tau and p-tau were normalized to GAPDH. SP tau was compared directly between IgG1 and 8B2 treated group. Unpaired t test in (B), (C), (D) and (F). Colocalization was quantified by Mander’s coefficiency. Scale bar, 50 μm.

Tauopathy mice exhibit gliosis, and IgG1 antibody subclass can elicit a strong microglia effector function. Therefore, we evaluated the glia status of JNPL3 mice after either 8B2 or control IgG1 treatment. The antibody 8B2 showed a strong colocalization with microglia marker Iba1, whereas the control IgG1 barely overlapped with Iba1 (Figure S2A-B). Moreover, 8B2 treatment significantly decreased Iba1 positive area compared to IgG1 control treated JNPL3 mice (Figure S2C). There was no difference in the colocalization of 8B2/GFAP and IgG1/GFAP (Figure S2D-E). However, mice treated with 8B2 exhibited less GFAP positive astrocytes compared to IgG1-treated mice (Figure S2F). In addition, the level of GFAP and Iba1 both significantly decreased in the forebrain lysates from mice treated with 8B2, compared to those treated with control IgG1 (Figure S2G-I). Overall, 8B2 treatment decreased microgliosis and astrogliosis in JNPL3 mice. Moreover, 8B2 was taken up by microglia that likely facilitated the degradation of pathological tau in these mice.

### Partially restored somatic Ca^2+^ activity in 6 month old JNPL3 mice acutely treated with 8B2

Considering the excellent target engagement of 8B2 antibody and its efficacy to reduce pathological tau, we tracked the Ca^2+^ activity in the same neuronal soma before and after 8B2 treatment, which was not performed in our previous study ^1^. In resting JNPL3 mice, Ca^2+^ transient frequency and total Ca^2+^ activity significantly increased after 8B2 treatment (Figure 8A and 8C). The fraction of hypoactive neurons decreased (38% vs 30%, before vs after, Figure S3A) and the fraction of hyperactive neurons increased (6% vs 12%, before vs after, Figure S3A). The peak amplitude of Ca^2+^ transients remained the same before and after 8B2 treatment (Figure 8B and S3B). In running JNPL3 mice, the frequency of Ca^2+^ transients and total Ca^2+^ activity were similar before and after 8B2 treatment (Figures 8D, S3A and 8F), but peak amplitude of Ca^2+^ transients significantly increased after 8B2 treatment (Figure 8E and S3B).

**Figure 8.**
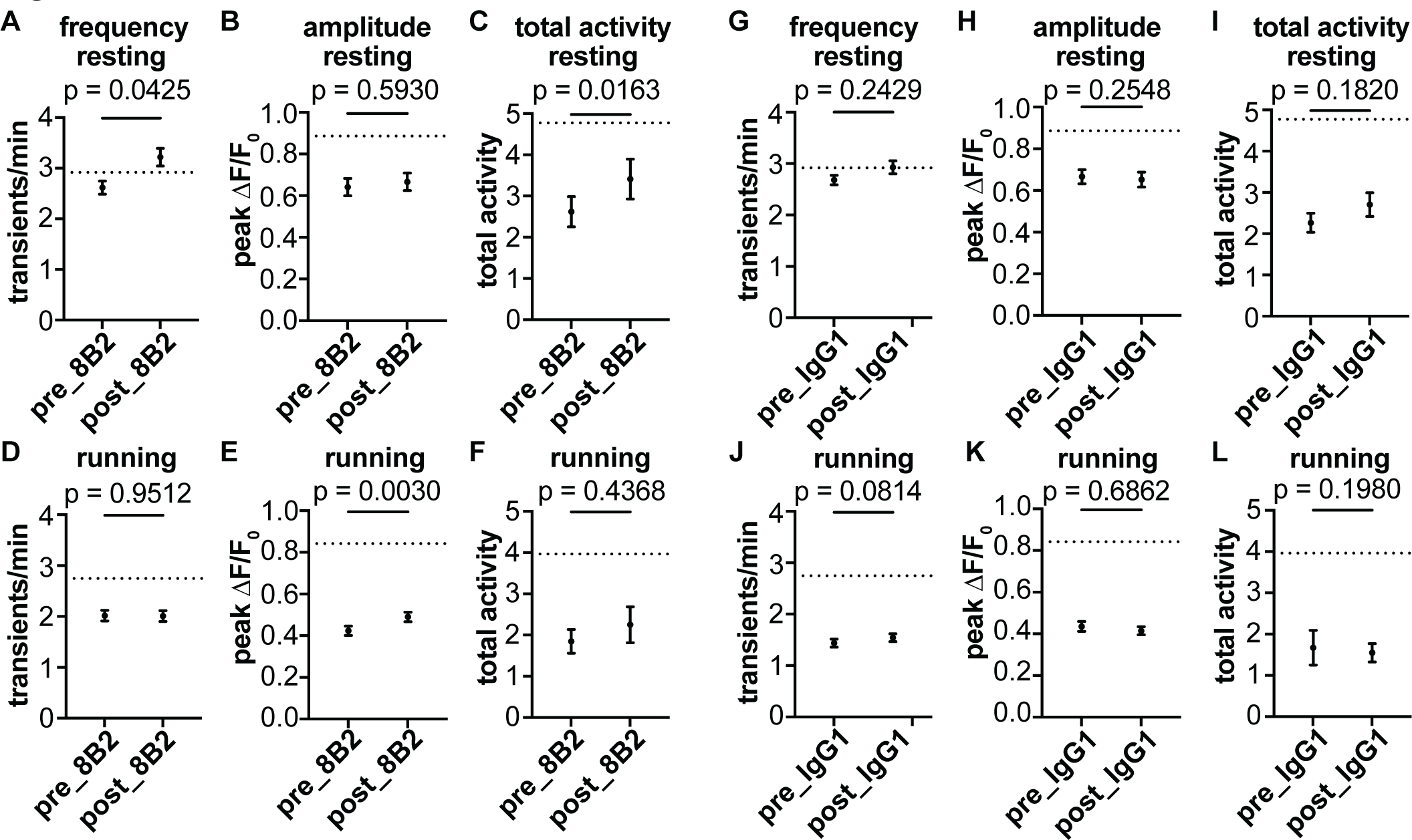
Abnormal Ca^2+^ activity is partially restored in L2/3 motor cortical neurons of JNPL3 mice after acute tau antibody 8B2 treatment. **(A - F)** Ca^2+^ activity frequency, peak amplitude, and total neuronal activity were analyzed in JNPL3 mice treated with 8B2 when animals were either resting or running on the treadmill. The Ca^2+^ activity in the same neuron before and after treatment were compared. In the resting condition, frequency of Ca^2+^ transients significantly increased to WT level (dotted line). Total activity of Ca^2+^ transients also significantly improved after 8B2 treatment. In the running condition, peak amplitude of Ca^2+^ transients significantly increased after 8B2 treatment. A total of 233 neurons from 6 JNPL3 mice were analyzed. **(G - F)** Ca^2+^ activity in JNPL3 mice treated with control IgG1. Same analyses were conducted as in (A - F). Control IgG1 had no effect to correct Ca^2+^ activity abnormalities. A total of 349 neurons from 7 JNPL3 mice were analyzed. Wilcoxon matched-pair signed rank test in (A-F).

Furthermore, we examined whether 8B2 treatment restored altered neuronal network activity during running in JNPL3 mice. Although there were trends of increase in activated neurons and decrease of suppressed neurons after acute 8B2 treatment, the changes did not reach statistical significance (Figure S3C-E). In addition, 8B2 treatment did not affect altered Ca^2+^ activity synchrony in either activated or suppressed neuronal population, respectively (Figure S3F).

To test if the functional improvement was specific to 8B2 treatment, we analyzed Ca^2+^ activity profiles in a cohort of JNPL3 mice treated by control IgG1. Two doses of control IgG1 treatments did not alter the frequency or amplitude of Ca^2+^ transients, nor total Ca^2+^activity in either resting or running JNPL3 mice (Figure 8G-L). Unlike 8B2 treatment, control IgG1 treatment increased the fractions of hypoactive and hyperactive neurons, and decreased the fraction of normal neurons under resting condition (Figure S4A). During running, the fractions of these three different types of neurons remained same before and after control IgG1 treatment (Figure S4A). Moreover, control IgG1 did not alter the pattern of neuronal responses to running (Figure S4B-D). In addition, we observed no change in Ca^2+^ activity synchrony among activated neurons or suppressed neurons during running after control IgG1 treatment (Figure S4E).

We also treated a cohort of WT mice with two doses of 8B2 to investigate if 8B2 treatment affected Ca^2+^ activity in healthy mice. No significant changes in frequency or amplitude of Ca^2+^ transients or total Ca^2+^ activity were observed in resting or running WT mice after 8B2 treatment (Figure S5A). In addition, the fractions of hypoactive, hyperactive and normal neurons remained similar before and after 8B2 treatment (Figure S5B).

Taken together, these data show that two doses of 8B2 treatment partially restored somatic Ca^2+^ activity profiles in motor cortical neurons in JNPL3 tauopathy mice under resting and running conditions. However, it did not restore the abnormal neuronal network activity in these mice. Nevertheless, this acute treatment paradigm and the assessment of neuronal functional readouts can be further optimized and utilized as a foundation to evaluate the therapeutic potential of tau antibodies in future studies.

## Discussion

In this study, we detected abnormal Ca^2+^ activity in motor cortical neurons of 6 month old awake and behaving JNPL3 mice, which were in their early stage of tauopathy. More pronounced changes of Ca^2+^ activity were observed in 6 month old cohorts compared to 12 month old cohorts when comparing JNPL3 mice and their age-matched WT controls. In resting 6 month old animals, a notable decrease in both peak amplitude and total activity of Ca^2+^ transients was observed in JNPL3 mice compared to WT mice. Moreover, running JNPL3 mice exhibited a more pronounced reduction of Ca^2+^ transient frequency, amplitude and total Ca^2+^ activity, compared to WT mice. In 12 month old animals, only peak amplitude of Ca^2+^ transients decreased significantly in resting or running JNPL3 mice compared to WT mice. The attenuation of Ca^2+^ activity in JNPL3 mice at 6 months of age did not further deteriorate at 12 months of age, although a significant decrease of total Ca^2+^ activity was observed in older WT mice compared to the younger WT mice. We further analyzed the Ca^2+^ activity profiles of the 6 month old cohorts in details. A large fraction of neurons was hypoactive in JNPL3 mice. Analyses of neuronal response to treadmill running revealed that JNPL3 mice had less activated neurons and more suppressed neurons compared to WT mice. Despite fewer activated neurons in JNPL3 mice, higher synchrony of Ca^2+^ activity was observed when animals were resting, and disrupted engagement of neuronal Ca^2+^ activity during running. Altered excitatory synaptic transmission and enhanced intrinsic neuronal excitability were also observed, which likely contributed to the decreased Ca^2+^ activity and disrupted neuronal synchrony in JNPL3 mice. Lastly, we attempted to improve neuronal and network function by acute tau antibody treatment. Two doses of acute tau antibody 8B2 treatment partially restored Ca^2+^ activity profiles but not network activity in JNPL3 mice. The 8B2 antibody was taken up by neurons and microglia and colocalized with total tau and phosphorylated tau. Acute 8B2 treatment decreased soluble phosphorylated tau, increased total insoluble tau and attenuated micro- and astrogliosis in JNPL3 mice.

A major observation in this study was decreased Ca^2+^ activity in L2/3 motor cortical neurons of 6 month old JNPL3 mice. This deficit was further exacerbated, compared to WT controls, when animals were running on a treadmill. Specifically, not only did a larger fraction of neurons become silent or hypofunctional, but the remaining active neurons also exhibited less Ca^2+^ activity in JNPL3 mice. The changes in Ca^2+^ transient amplitude and total Ca^2+^ activity were similar to our prior analyses of older 10-12 month old JNPL3 mice, while changes in the frequency of Ca^2+^ transients differed from that prior report ^1^. This suggests persistent but evolving changes in neuronal activity during the progression of tau pathology. Reduced neuronal firing rates or Ca^2+^ transient frequency has been reported in other mouse models at their early stage of tauopathy ^5,6^. However, these two studies did not explore the changes in the Ca^2+^ activity amplitude nor the total Ca^2+^ activity, especially when animals were awake and performing a task like running.

Previous in vivo two-photon study or in vivo electrophysiological recordings by others were performed in tauopathy mice that were anesthetized or awake and resting ^5,6^. In our prior and current studies, the mice were performing a running task, which is a relatively strong stimulus ^1^. We previously observed exacerbated Ca^2+^ abnormalities in running vs resting JNPL3 tauopathy mice at 10-12 months of age, compared to their age-matched WT controls ^1^. In the current study, we adopted the same experimental procedure. In addition to performing similar analyses as we did before on both young (6 months) and old (12 months) animals, we also analyzed the Ca^2+^ activity profiles and neuronal network activity in the younger cohorts. We are not aware of prior such analyses in tauopathy mice. We found varying responses of L2/3 motor cortical neurons to running, with some neurons being activated whereas others were suppressed. In JNPL3 mice, the fraction of activated neurons significantly decreased, while the fraction of suppressed neurons significantly increased. This further revealed reduced neuronal activity in running JNPL3 mice, and the proper neuronal responses to running behavior was disrupted in these animals.

Even though neurons in tauopathy mice showed reduced activity, the synchrony of basal neuronal Ca^2+^ activity significantly increased in resting 6 month old JNPL3 mice, which was not examined in our previous study in older animals ^1^. Increased neuronal synchrony has been observed in hyperactive neurons in an AD mouse model with Aβ deposits ^23^. However, neurons in Aβ mouse models usually exhibit hyperactivity ^6,20,24^, which is in contrast to the hypoactivity in tauopathy models reported by us and others ^1,5,6,25,26^. Nevertheless, excessive neuronal synchrony can lead to elliptical like discharge in AD patients and mouse models ^13^. Considering these data with our findings, we hypothesize that pathological tau may synergize with Aβ to increase synchronization of basal neuronal activity. When mice were running, neuronal Ca^2+^ activity synchrony also increased in JNPL3 mice. However, neurons with different responses to running showed diverging changes. Those that were activated during running were more synchronized, while those with suppressed activity were less synchronized. These data indicate that JNPL3 mice had disrupted neuronal circuitry engagement in L2/3 motor cortex during running.

We further investigated the underlying mechanisms causing reduced neuronal Ca^2+^ activity and altered network activity in this study. Although the Ca^2+^ activity detected by GCaMP6 indicator cannot fully depict action potential firing patterns, decreased somatic Ca^2+^ transient frequency in running JNPL3 mice may link to suppressed intrinsic excitability or synaptic transmission. Our previous study in the 10-12 month old cohort of the same model observed significantly decreased synaptophysin and AMPA receptors in JNPL3 brain lysates ^1^. Those data indicate impaired excitatory synaptic function associated with tau pathology. Other have shown that pathological tau accumulated at presynaptic terminals leads to synaptic vesicle immobility, loss of synaptic boutons and deficits of synaptic plasticity in entorhinal cortex and hippocampus ^27–31^ In post-synaptic density, tau has been shown to be involved in AMPA receptor trafficking and indirectly modulate NMDA receptor function ^32–38^. In addition, mutant tau has been reported to obstruct AMPAR trafficking during LTP or cause NMDR-mediated excitotoxity ^32–35,37,38^. Further studies on how tau-mediated synaptic dysfunction contributes to altered neuronal network in behaving animals are warranted.

In a previous study, accumulation of hyperphosphorylated tau reduced excitability of hippocampal neurons in young 3xTg mice ^39^. However, we did not observe a decrease of intrinsic excitability in L2/3 motor cortical neurons of our tauopathy mice. Instead, both FS inhibitory neurons and non-fasting-spiking neurons exhibited enhanced excitability in 6 month old JNPL3 mice. This difference is possibly due to different brain regions and animal models used in ours vs that study. In our mice, the excitability of FS neurons was slightly enhanced in JNPL3 mice. However, mutant tau did not change input resistance, capacitance or rheobase current of FS neurons. Therefore, the enhanced firing ability of FS neurons was likely associated with increased excitatory inputs onto these cells, specifically increased sEPSC frequency. Consequently, increased FS neurons firing might contribute to enhanced synchrony of neuronal activity given the roles of FS interneurons in network oscillations ^40^. In contrast, non-fasting-spiking neurons in our JNPL3 mice exhibited greater increase of excitability. The mutant neurons displayed larger input resistance, likely due to a decrease of basal repolarization conductance. As a result, these neurons had depolarized resting membrane potential, and needed lower rheobase current to fire action potentials. Enhanced excitability in non-fasting-spiking neurons may be a compensatory response to decreased excitatory synaptic transmission onto them, indicated by a decrease in sEPSC frequency through homeostatic mechanisms ^22^. Nevertheless, these disproportional changes of excitability in FS inhibitory neurons and non-fasting-spiking neurons (presumably pyramidal excitatory neurons) might contribute to imbalance between excitatory and inhibitory activity (E/I imbalance) and compromise local network activity in JNPL3 mice. Further studies on how pathological tau alters firing properties in different neuronal subtypes are needed. Given that E/I balance modulated neuronal firing synchrony in physiological conditions according to previous studies ^12,13,41^, further exploration of how altered E/I balance disrupted neuronal population activity in pathological conditions like tauopathy is also needed to clarify circuit mechanisms of network dysfunction in behaving animals.

Lastly in our study, we evaluated the therapeutic efficacy of a tau antibody 8B2 in the 6 month old cohort of JNPL3 mice. We used the same acute treatment paradigm as in our previous report on tau antibody 4E6 in 10-12 month old JNPL3 mice ^1^. The 8B2 antibody targets the pS404 epitope of tau like 4E6, but their affinities and exact binding site differ substantially ^17^. Different from our analyses in Wu et al, we aligned images obtained before and after antibody treatment, and tracked the Ca^2+^ activity of individual neurons across different days, which enhances the power of the analyses. Two acute doses of 8B2 restored some of the altered Ca^2+^ activity in JNPL3 mice, namely Ca^2+^ transient frequency and total Ca^2+^ activity in resting animals, and Ca^2+^ transient amplitude in running animals. However, it failed to rescue altered population patterns of Ca^2+^ activity and the network engagements to running in JNPL3 mice. Two-doses of 8B2 treatment decreased soluble phosphorylated tau and increased insoluble tau. Additionally, 8B2 antibody was efficiently taken up by neurons and microglia and colocalized with tau, indicating target engagement. Furthermore, treatment with 8B2 antibody significantly decreased microgliosis and astrogliosis in JNPL3 mice. Altogether, these data support functional benefits of acute antibody 8B2 treatment in 6 month old JNPL3 mice, supporting chronic studies on this antibody that should provide greater efficacy.

### Summary

In the JNPL3 tauopathy mouse model, reduction of neuronal Ca^2+^ activity was observed in motor cortex of awake and behaving animals at their early stage of tauopathy that was exacerbated under running condition, compared to age-matched WT controls. This neuronal deficit did not further deteriorate from 6 to 12 months of age. Furthermore, the tau pathology affected local neuronal circuitry, resulting in increased synchrony of basal Ca^2+^ activity and altered running-related neuronal responses in the L2/3 of primary motor cortex. Underlying mechanisms may link to impairments of excitatory synaptic transmission and altered E/I balance in this region resulting from accumulation of pathological tau. Tau antibody treatment cleared soluble phospho-tau, increased insoluble total tau and ameliorated gliosis while partially restoring abnormal Ca^2+^ activity profile, highlighting the importance of functional assessment when evaluating the therapeutic efficacy of anti-tau therapies.

## Materials and Methods

### Animals

Homozygous transgenic JNPL3 mice expressing human 0N4R tau with P301L mutation (Taconic) and age-matched wild-type (WT) mice of the same strain background at 6 and 12 months of age were used in this study. We chose females as in our previous study because they have more tau pathology than males ^1^. All animals were housed at NYU Grossman School of Medicine animal facilities. All procedures were approved by the Institutional Animal Care and Use Committee (IACUC) of the university, and are in accordance with NIH Guidelines, which meet or exceed the ARRIVE guidelines.

### Surgery

Cranial window surgery was performed as we have described previously ^1^. Briefly, the mouse was placed in a stereotaxic frame with a heated pad under isoflurane anesthesia. A round opening of 3 mm in diameter was drilled into the skull over the right primary motor cortex. Subsequently, a total of 0.6 µL of AAV5-Syn-GCaMP6s virus (1.8 × 10^13^ genome copies per mL, Addgene) was slowly injected by Nanoject III Nanoliter Injector into layer 2/3 motor cortex (1.5 mm anterior from bregma, 1.5 mm lateral from midline) to fluorescently label the neurons. Then the skull opening was covered by a round coverslip. Dental cement was used to seal the edges of the cover glass and embed a head holder composed of two parallel micro-metal bars. Mice were individually housed and allowed to recover for four weeks before two-photon imaging.

### In vivo two-photon microscopy

A custom-built free-floating treadmill was placed under the microscope. Mice were head-fixed and allowed to move their forelimbs on the treadmill to perform a running task. Mice were first placed on the treadmill to record the Ca^2+^ activity in a quiet restful state (referred to as resting) for 100 s. Then, the treadmill was turned on and the mice were forced to run at a speed of 1.67 cm/s (referred to as running). When the motor was turned on, the belt speed of the treadmill gradually increased from 0 cm/s to 1.67 cm/s within 2 s. Each mouse ran five trials with a short resting period (25 s) in between trials. Each running trial lasted for 100 s.

Two-photon imaging was collected with an Olympus Fluoview 1000 two-photon system (920 nm) equipped with a Ti:Sapphire laser (MaiTai DeepSee, Spectra Physics). Ca^2+^ signals were recorded at 2 Hz using a 25X objective (NA 1.05), with a frame size of 320 × 256 pixels with a 1X digital zoom or 256 × 256 pixels with a 2X digital zoom. Images were taken at the depth of 200-300 µm below the pial surface. The same focal plane with same neurons was imaged again after antibody treatment. Images were acquired by Fluoview software and analyzed post hoc using ImageJ, MATLAB, Python and R/RStudio.

### Image analysis

Individual images were first motion-corrected in Fiji Image J using the plugin Image Stabilizer (https://imagej.net/Fiji). Subsequently, all images from different trials of the same animals were aligned by the motion correction function of Using_the_Acquisition2P_Class.m in MATLAB (https://github.com/HarveyLab/Acquisition2P_class/tree/master). Once the trial alignments were finished, all the images from the same animals were concatenated chronologically into a single image in Image J. This image was then analyzed using the Mesmerize package ^15^. Regions of interest (ROI) corresponding to neuronal somas and the time-dependent component of relative fluorescence change df/f within each ROI were extracted using CaImAn toolbox in Mesmerize. The CNMF method in CaImAn toolbox allows eliminating signal contamination from background and nearby neuropils ^16^. Subsequently, df/f’s were exported from Mesmerize for further analyses.

The frequency of Ca^2+^ transients, amplitude of transients and total Ca^2+^ activity was analyzed in a custom-written R script ^1^. Briefly, a time window spanning 15% of the trace length was slid across the trace, and the trace with minimal standard deviation (SD) within a window was selected as baseline. Threshold was set at the average of baseline plus five times its SD. The frequency of Ca^2+^ transients was calculated by counting the number of Ca^2+^ transients per minute for each soma. The peak amplitude was defined as the highest value of the Ca^2+^ transients in each trace. The mean amplitude was the average of all Ca^2+^ transients in a given trace. Finally, the total Ca^2+^ activity was quantified by calculating the area under the curve of Ca^2+^ transients per minute.

To analyze running-related responses, Ca^2+^ activity (df/f) of each neuron was averaged across five trials. The averaged trace of each neuron was then aligned to the time when the treadmill started. Neurons were categorized into different groups by comparing Ca^2+^ activity (df/f) before and after the start of the treadmill (stationary vs moving). Specifically, the baseline activity of each neuron was calculated by averaging its activity during resting and was subtracted from the averaged df/f during running. After the treadmill started, if the baseline-subtracted activity deviated more than three times the SD of baseline for at least 3 consecutive frames (equivalent to 1.5 s), the neuron was identified as one that responded to running. The neuron was further categorized as ‘activated’ or ‘suppressed’ if the df/f was above or below the baseline, respectively. When deviations occurred in both directions, the neuron was categorized as mixed-response neurons. The rest of the neurons show no change between resting vs running.

Pearson correlation coefficiency of calcium activity (df/f) was calculated between each pair of neurons for each field of view. The mean correlation value of a given cell was averaged from all the correlation coefficiencies between that cell and all other simultaneously recorded cells within the same field of view. The correlation matrices were calculated separately when animals were either resting or running, and averaged from five trials.

### In vitro brain slice recordings

WT or JNPL3 female mice (6-7 month old) were first anesthetized with isoflurane and then decapitated. Brains were quickly removed and immersed in ice-cold oxygenated cutting solution containing (in mM) 87 NaCl, 2.5 KCl, 1.25 NaH_2_PO_4_, 25 NaHCO_3_, 10 glucose, 75 sucrose, 1.3 ascorbic acid, 7 MgCl_2_, 0.5 CaCl_2_. Coronal slices (350 um) were cut with a vibrating microtome (Leica), and immediately transferred to a chamber with oxygenated (95% O_2_/5% CO_2_) artificial cerebrospinal fluid (ACSF) containing (in mM) 124 NaCl, 2.5 KCl, 1.25 NaH_2_PO_4_, 26 NaHCO_3_, 10 glucose, 1.5 MgCl_2_, 2.5 CaCl_2_ at 35 °C for 20 min. The chamber with slices was then transferred to room temperature to allow the slices to recover for at least 1 h. For recording, slices were placed in a recording chamber perfused with oxygenated ACSF that was heated to 30 °C. The recording chamber was held under an Olympus microscope equipped with infrared differential interference optics. Chemicals were all purchased from Sigma-Aldrich unless stated otherwise.

Whole-cell recording was performed on individual neurons within layer 2-3 of primary motor cortex. Patch pipettes were pulled from borosilicate glass (World Precision Instruments, 140N-15) with an access resistance of 3-5 MΩ. These pipettes were filled with internal solution containing (in mM) 127 K-gluconate, 8 KCl, 10 phosphocreatine, 10 HEPES, 4 Mg-ATP, 0.3 Na-GTP (pH7.3, 280 mOsm). After membrane breaking to form whole cell configuration, the resting membrane potential was measured for each neuron. Subsequently, action potentials (AP) were stimulated by injecting a series of depolarization current steps from a holding potential of -70 mV. Data were recorded at a sampling rate of 20 kHz and filtered at 2 Hz by a 8-pole Bessel filter using a MultiClamp 200 amplifier (Molecular Devices). Cells with resting membrane potential more depolarized than -50 mV and access resistance exceeding 25 MΩ were excluded from further analyses. Fast-spiking (FS) or non-FS cells were identified based on their AP firing patterns. AP waveforms were analyzed by custom-written Matlab scripts.

To record spontaneous excitatory postsynaptic current, the internal solution to record AP was used. The cell types were first identified by AP firing patterns and then followed by sEPSC recording. The membrane potential was held at -70 mV which was at the reversal potential of Cl^-^. Data were analyzed in Clampfit (Molecular Devices).

### Antibodies and intravenous injections

Mouse monoclonal tau antibody 8B2 IgG1κ was generated by GenScript as previously described by immunizing BALB/c mice with a peptide encompassing to pSer396/404 region of the tau protein that was conjugated to keyhole limpet hemocyanin via a cysteine residue (cTDHGAEIVYK(pS)PVVSGDT(pS)PRHL) ^17^. Hybridoma fusions were screened by ELISA, and 8B2 was selected with other antibodies based on their binding to the phospho-tau peptide immunogen. We have previously reported on its crystal structure and binding characteristics ^17^. IgG1κ (referred as IgG1, eBioscience, 16–4714) was used as a control of the same subclass. IgG1 or 8B2 antibody (100 µg per injection) were injected intravenously into JNPL3 mice on Day 1 and Day 4 after the initial image session (Day 0). The dose of each injection equals 3 mg/kg antibody. A subset of mice was injected intravenously with antibodies labeled with VivoTag 680XL to assess target engagement.

### Tissue processing

After imaging, the brains were collected for downstream analyses. Briefly, mice were perfused with PBS before brain extraction. The right hemispheres were fixed in 4% paraformaldehyde at 4°C overnight, and then transferred into 2% DMSO and 20% glycerol in phosphate buffer for long-term storage at 4°C until cryosection. Brain slices were sectioned coronally at 40 μm for immunohistology. The left brain hemispheres were flash frozen on dry ice and stored at -80°C for subsequent protein assays and biochemical analyses.

### Western blots

The left brain hemispheres were homogenized in ice-cold modified RIPA buffer (50 mM Tris-HCl, 150 mM NaCl, 1 mM EDTA, 1% Nonidet P-40, 0.25% sodium deoxycholate, pH 7.4) with protease cocktail (cOmplete, Roche) and phosphatase inhibitor cocktail (1 mM NaF, 1 mM Na_3_VO_4_, 1 nM PMSF). The brain homogenates were centrifuged at 20,000xg for 1 h at 4°C. The resulting supernatants were collected as low speed supernatant (LSS). To isolate the sarkosyl insoluble fraction, 2 mg total protein from LSS was diluted in RIPA buffer with 1% sarkosyl, and then centrifuged at 100,000xg for 1.5 h at 20°C. The pellet was washed by RIPA buffer with 1% sarkosyl and centrifuged again at 100,000xg for 1 h at 20°C. The sarkosyl pellet (SP) was air dried for 30 min, and then dissolved in 50 μl of 5X sample buffer (62.5 mM Tris-HCl, 10% glycerol, 5% β-mercaptoethanol, 2.3% SDS, 1 mM EDTA and 1 mM EGTA). The LSS was eluted with 1X sample buffer. All samples were boiled for 10 min, followed by electrophoresis on 12% SDS-PAGE gel and transferred onto nitrocellulose membrane. The blots were blocked by 5% milk in TBST (Tris buffered saline with 1% TritonX-100). Primary antibodies, including PHF1 (1:1000, gift from Peter Davies), CP27 (1:500, gift from Peter Davies), pT231 (1:2000, Thermo Fisher Scientific, 1H6L6), GAPDH (1:5000, Cell Signaling Technology, D16H11), GFAP (1:2000, Bioss, bs0119R), Iba1 (1:2000, Fujifilm Wako, 019-19741) were diluted in Superblock® (Thermo Fisher Scientific), and incubated with the membrane at 4°C overnight. The following day, the membrane was incubated with diluted (1:10,000) IRDye 800CW goat anti-rabbit or IRDye® 680RD goat anti-mouse secondary antibodies (LICOR Biosciences). Blots were then detected by Odyssey® CLx imaging system (LICOR Biosciences).

### Immunohistochemistry

Brain sections were washed in PBS, followed by permeabilization and blocking in PBS with 5% bovine serum albumin, 2% normal goat serum and 0.3% Triton-X-100 at room temperature for 3 h. Subsequently, the sections were incubated with diluted primary antibodies in PBS with 5% bovine serum albumin and 1% normal goat serum at 4°C overnight. The dilution of the primary antibodies was as follows: PHF1 (1:100, gift from Peter Davies), CP27 (1:100, gift from Peter Davies), NeuN (1:200, Cell Signaling Technologies, D4G4O), GFAP (1:200, Bioss, bs0119R), Iba1 (1:200, Fujifilm Wako, 019-19741). On the following day, the brain sections were incubated with diluted secondary antibodies (1:1000) goat anti-mouse Alexa Fluor 568 or goat anti-rabbit Alexa Fluor 568 at room temperature for 3 h. After washing, the sections were mounted in ProLong Gold antifade reagent (ThermoFisher, P10144), and imaged by an LSM 700 Zeiss confocal laser scanning microscope. Images were processed by Fiji ImageJ software.

### Statistics

Statistics were performed using GraphPad Prism 9. Data are presented as mean ± standard error of the mean (SEM). Prior to applying statistical comparisons, data distributions were assessed by Shapiro-Wilk normality test. For normally distributed data, paired or unpaired t-tests were used, while non-normally distributed data were analyzed using the Mann-Whitney U test or Kolmogorov-Smirnov test. The specific tests used for each figure are indicated in the corresponding figure legend and summarized in Table 1. All comparisons were two-tailed, and a significance level was set to p ≤ 0.05.

**Table 3.** Ca^2+^ transient frequency, spike number per second; Ca^2+^ transient amplitude, peak ΔF/F_0_; integrated Ca^2+^ transient AUC, Ca^2+^ transient AUC per minute; Ca^2+^ activity correlations, Pearson’s correlation of ΔF/F_0_ over time between neuronal pairs in a given field of view.

|  | Ca<sup>2+</sup> activity profiles | 6 months |  | 12 months |  |
| --- | --- | --- | --- | --- | --- |
|  |  | JNPL3 vs WT |  | JNPL3 vs WT |  |
|  |  | resting | running | resting | running |
| Animal level | Ca<sup>2+</sup> transient frequency | ↔ | ↓ | ↔ | ↔ |
|  | Ca<sup>2+</sup> transient amplitude | ↓ | ↓ | ↓ | ↓ |
|  | integrated Ca<sup>2+</sup> transient AUC | ↓ | ↓ | ↔ | ↔ |
|  | Ca<sup>2+</sup> activity correlations | ↑ | ↔ | NA | NA |
| Neuronal level | fraction of hypoactive neurons | ↔ | ↑ | NA | NA |
|  | Ca<sup>2+</sup> transient frequency of active neurons | ↔ | ↓ | NA | NA |
|  | Ca<sup>2+</sup> transient amplitude of active neurons | ↓ | ↓ | NA | NA |
|  | integrated Ca<sup>2+</sup> transient AUC of active neurons | ↓ | ↓ | NA | NA |
|  | Ca<sup>2+</sup> activity correlations | ↑ | ↑ | NA | NA |
|  | fraction of activated neurons | NA | ↓ | NA | NA |
|  | fraction of suppressed neurons | NA | ↑ | NA | NA |
|  | Ca<sup>2+</sup> activity correlations of activated neurons | NA | ↑ | NA | NA |
|  | Ca<sup>2+</sup> activity correlations of suppressed neurons | NA | ↓ | NA | NA |

## Acknowledgements

This work was supported in part by NIH grants R21 AG059391, R01 AG032611, R01 NS077239, and R01 NS120488 (E.M.S.), and the Alzheimer’s Association Research Fellowships [AARF-22-926735 (C.J.), AARF-22-924783 (Y.J.), AARFD-22-926379 (A.M-A)].].

We also thank the technical support from NYU Langone Health’s Ion Laboratory (RRID: SCR_021754) for electrophysiology studies.

## Author contributions

C.J. and E.M.S. conceived the project and wrote the article. C.J. and X.Y. performed the experiments and related analyses. M.E. analyzed some of the electrophysiological data. Y.J. prepared the labelled 8B2 antibody and control IgG. A.M.T. performed immunofluorescent staining of the mouse brains. S.C.S. provided guidance on electrophysiological recording and data analyses. A.M.A, Q.W, Y.Z. and W.G. provided guidance on two-photon calcium imaging experiments. Y.L. maintained the animal colonies. All authors had the opportunity to edit the article. E.M.S. supervised the project.

## Data and materials availability

All data needed to evaluate the conclusions in the paper are present in the paper and/or the Supplemental Materials.

## Ethics approval and consent to participate

All procedures were approved by the Institutional Animal Care and Use Committee (IACUC) of the New York University Grossman School of Medicine, and are in accordance with NIH Guidelines, which meet or exceed the ARRIVE guidelines. The consent to participate is not applicable.

## Figure Legends

**Supplemental Figure 1.**
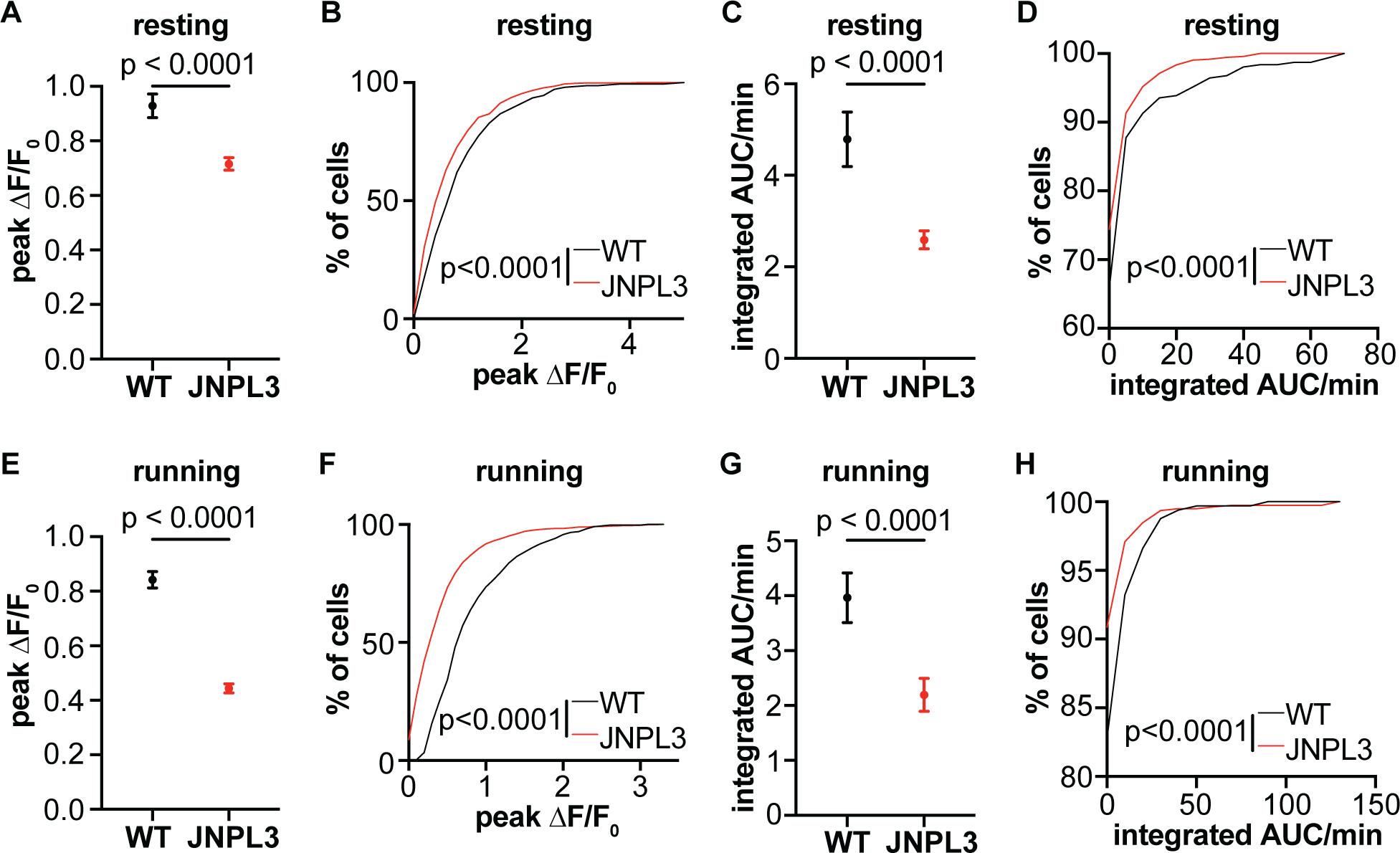
Decreased Ca^2+^ activity in L2/3 motor cortical neurons of 6-month-old JNPL3 mice. **(A)** The maximum Ca^2+^ transient amplitude in active neurons (transients/min > 0) decreased in resting JNPL3 mice compared to WT mice. **(B)** Cumulative frequency distribution of (A). **(C)** The total activity (integrated Ca^2+^ activity per min) in active neurons in resting JNPL3 mice, which decreased compared to WT mice. **(D)** Cumulative frequency distribution of (B). **(E)** The maximum Ca^2+^ transient amplitude in active neurons decreased in running JNPL3 mice. **(F)** Cumulative frequency distribution of (E). **(G)** The total activity (integrated Ca^2+^ activity per min) in active neurons in running JNPL3 mice, which decreased compared to WT mice. **(H)** Cumulative frequency distribution of (G). In resting condition, a total of 301 neurons from WT animals and 733 neurons from JNPL3 mice were analyzed. Under running condition, a total of 325 neurons from WT animals and 802 neurons from JNPL3 mice were analyzed. Mann-Whitney U test in (A), (C), (E) and (G). Kolmogorov-Smirnov test in (B), (D), (F) and (H).

**Supplemental Figure 2.**
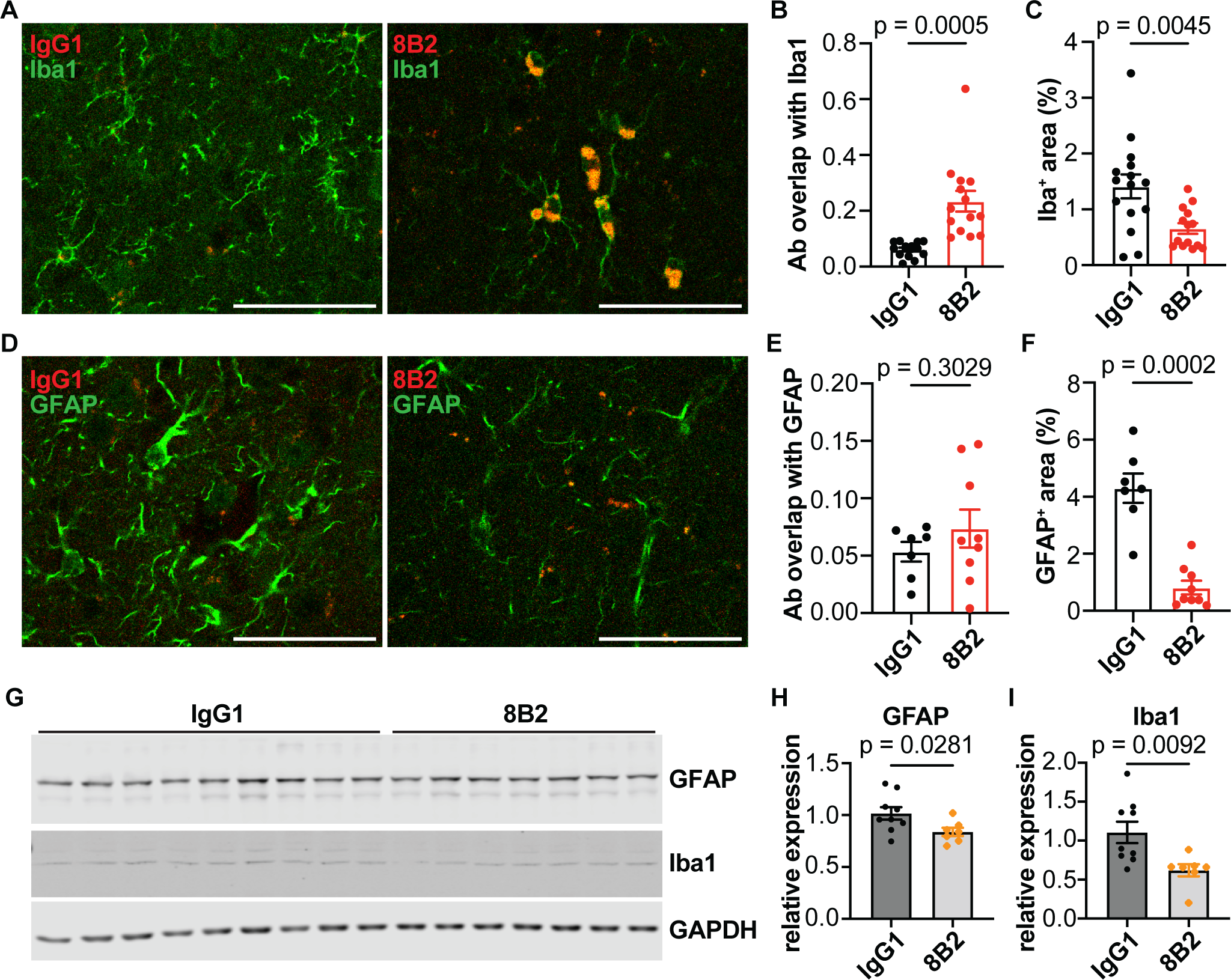
Attenuation of gliosis and accumulation of tau antibody 8B2 in microglia of the motor cortexes in 6 month old JNPL3 mice. These analyses were conducted on the same brains that were analyzed in Figure 7. **(A)** Coronal brain sections were stained with microglia marker Iba1. Colocalization between Iba1 and 8B2 was observed. **(B)** Overlapping signals between 8B2 and Iba1 were significantly more than those between IgG1 and Iba1. **(C)** Mice treated with 8B2 had less Iba1 reactive area than mice treated with control IgG1. **(D)** 8B2 or IgG1 had very limited overlap with astrocyte marker GFAP. **(E)** GFAP and 8B2 colocalization did not differ from GFAP and IgG1 colocalization. **(F)** Mice treated with 8B2 had less GFAP reactive area than IgG1 control treated mice. (A) - (F) showed a representative image from a mouse treated with either 8B2 or IgG1. **(G)** Western blots of GFAP and Iba1 in all animals. A total of 9 JNPL3 mice received IgG1 injections and 7 JNPL3 mice were injected with 8B2. **(H)** Quantification of GFAP expression normalized to GAPDH levels. Significant decrease of GFAP was observed in JNPL3 mice. **(I)** Quantification of Iba1 expression normalized to GAPDH levels. Significant decrease of Iba1 was observed in JNPL3 mice, too. Unpaired t test in (B), (C), (E), (F), (H), Colocalization was assessed by Mander’s coefficiency. Scale bar, 50 μm.

**Supplemental Figure 3.**
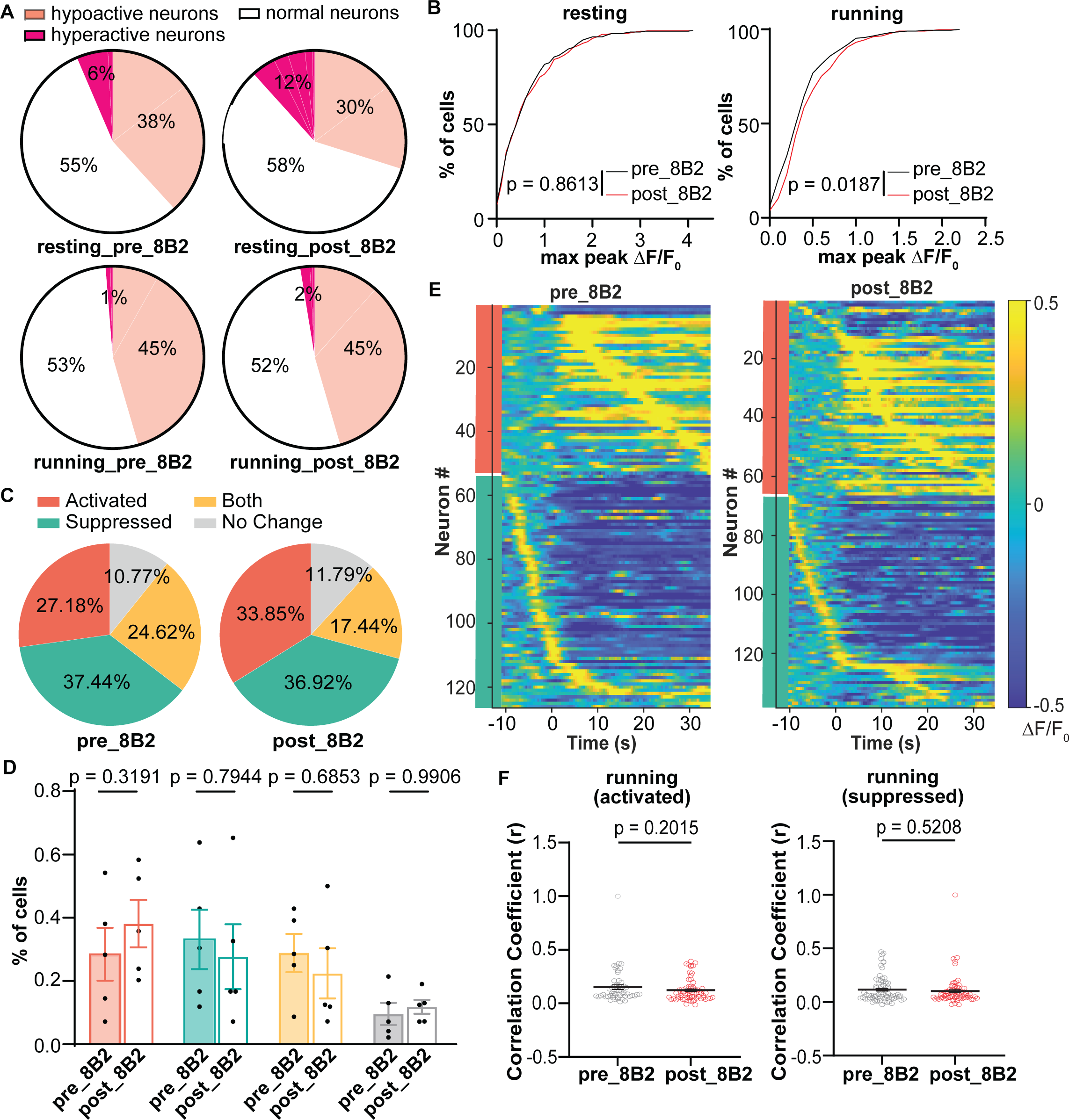
8B2 treatment failed to correct running-related neuronal Ca^2+^ activity abnormalities in L2/3 motor cortical neurons of JNPL3 mice. **(A)** The fractions of hypoactive neurons decreased in resting JNPL3 mice but remains unchanged in running mice after 8B2 treatment. **(B)** Cumulative frequency distribution of the maximal Ca^2+^ transient amplitude in neurons before and after 8B2 treatment. The distribution of maximal Ca^2+^ transient amplitude significantly changed after 8B2 treatment. **(C)** Fractions of neurons (195 neurons) categorized by their responses to running before and after 8B2 treatment. **(D)** Fractions of neurons with each response to running in JNPL3 mice (5 animals) before and after 8B2 treatment. Each dot represents an animal. **(E)** Activity patterns of neurons which were either activated or suppressed during running before and after 8B2 treatment. **(F)** Correlation index of activated or suppressed neurons before and after the 8B2 treatment. Each dot represented the correlation index of a given neuron. Kolmogorov-Smirnov test in (B). Multiple unpaired t test in (D). Unpaired t test in (F).

**Supplemental Figure 4.**
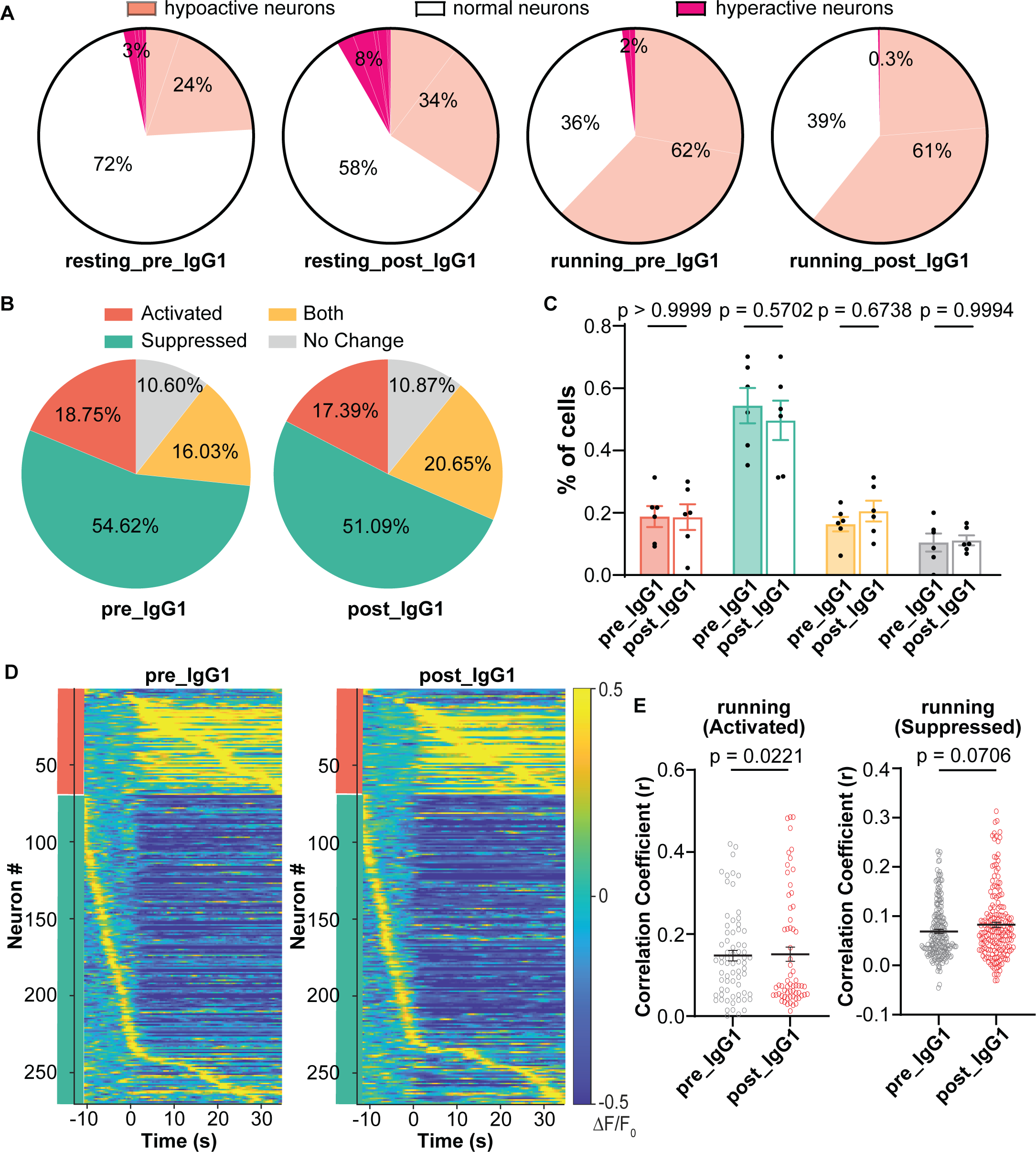
No change or exacerbated abnormal Ca^2+^ activity in L2/3 motor cortical neurons of JNPL3 mice treated with IgG1 controls. **(A)** The fractions of hypoactive neurons increased in resting JNPL3 mice or no change in running JNPL3 mice after IgG1 treatment. **(B)** Ratios of neurons (368 neurons) categorized by their responses to running before and after IgG1 treatment. **(C)** Fractions of neurons with each response to running in JNPL3 mice (6 animals) before and after IgG1 treatment. Each dot represents an animal. **(D)** Activity patterns of neurons with either activated or suppressed response to running before and after IgG1 treatment. **(E)** Correlation index of activated or suppressed neurons before and after IgG1 treatment. Each dot represented the correlation index of a given neuron. Multiple unpaired t test in (C). Unpaired t tests in (E).

**Supplemental Figure 5.**
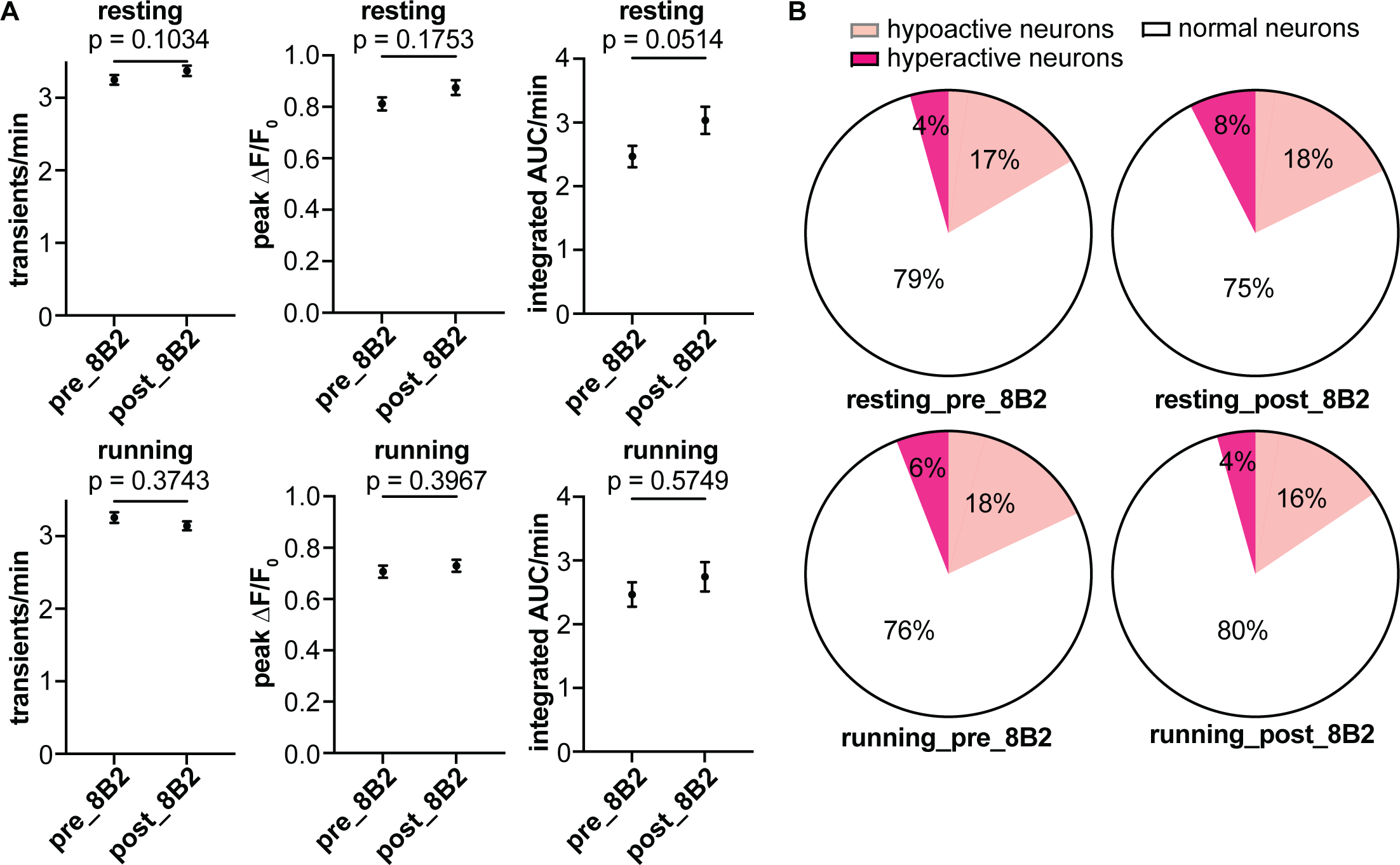
Somatic Ca^2+^ activity in L2/3 motor cortex before and after 8B2 injection in 12-month-old WT mice. **(A)** Ca^2+^ activity frequency, amplitude, and total neuronal were analyzed in 12-month-old WT mice when animals were either resting or running on the treadmill. No significant change of Ca^2+^ activity was observed in WT neurons before and after 8B2 treatment. **(B)** The fractions of hypoactive neurons remained unchanged in resting or running WT mice before and after 8B2 treatment. A total of 760 neurons from 5 WT mice were included in the analyses. Wilcoxon matched-pairs signed rank test in (A).

**Supplemental Figure 6.**
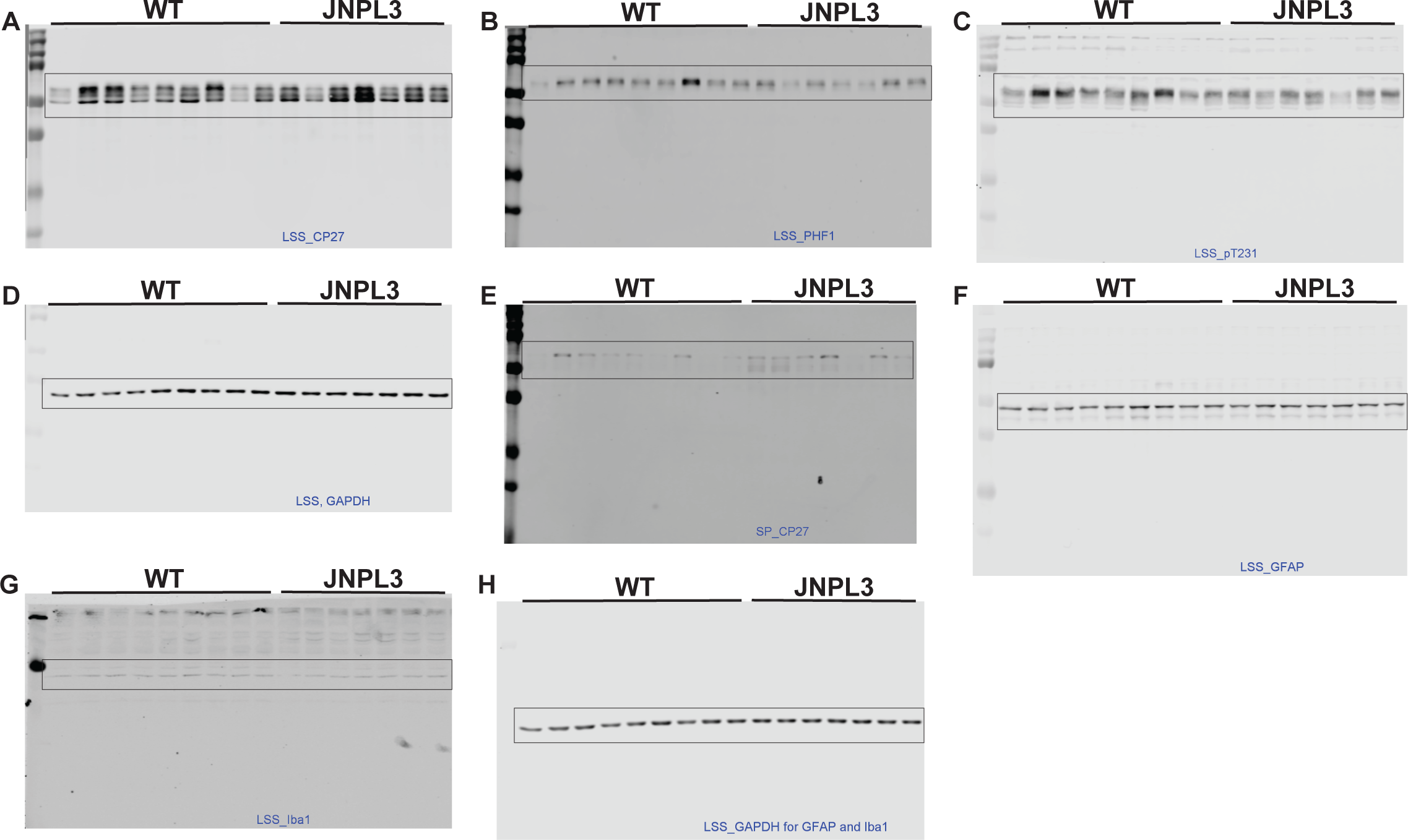
Full pictures of western blots. **(A) – (E)** were used in Figure 7C. **(F) – (H)** were used in Figure S2G. Boxes denoted the regions that were used in Figure 7C and Figure S2G.

## References

1. Wu Q, Bai Y, Li W, et al. Increased neuronal activity in motor cortex reveals prominent calcium dyshomeostasis in tauopathy mice. Neurobiol Dis. 2021;147. doi:10.1016/j.nbd.2020.105165

2. Scheltens P, De Strooper B, Kivipelto M, et al. Alzheimer’s disease. Lancet. 2021;397(10284):1577-1590. doi:10.1016/S0140-6736(20)32205-4

3. Robbins M, Clayton E, Kaminski Schierle GS. Synaptic tau: A pathological or physiological phenomenon? Acta Neuropathol Commun. 2021;9(1):149. doi:10.1186/s40478-021-01246-y

4. Wang Y, Mandelkow E. Tau in physiology and pathology. Nat Rev Neurosci. 2016;17(1):5-21. doi:10.1038/nrn.2015.1

5. Marinković P, Blumenstock S, Goltstein PM, et al. In vivo imaging reveals reduced activity of neuronal circuits in a mouse tauopathy model. Brain. 2019;142(4):1051–1062. doi:10.1093/brain/awz035

6. Busche MA, Wegmann S, Dujardin S, et al. Tau impairs neural circuits, dominating amyloid-β effects, in Alzheimer models in vivo. Nat Neurosci. 2019;22(1):57–64. doi:10.1038/s41593-018-0289-8

7. Lewis J, McGowan E, Rockwood J, et al. Neurofibrillary tangles, amyotrophy and progressive motor disturbance in mice expressing mutant (P301L) tau protein. Nat Genet. 2000;25(4):402–405. doi:10.1038/78078

8. Adler A, Zhao R, Shin ME, Yasuda R, Gan WB. Somatostatin-Expressing Interneurons Enable and Maintain Learning-Dependent Sequential Activation of Pyramidal Neurons. Neuron. 2019;102(1):202–216.e7. doi:10.1016/j.neuron.2019.01.036

9. Peters AJ, Chen SX, Komiyama T. Emergence of reproducible spatiotemporal activity during motor learning. Nature. 2014;510(7504):263–267. doi:10.1038/nature13235

10. Papale AE, Hooks BM. Circuit changes in motor cortex during motor skill learning. Neuroscience. 2018;368:283–297. doi:10.1016/j.neuroscience.2017.09.010

11. Buzsáki G, Watson BO. Brain rhythms and neural syntax: implications for efficient coding of cognitive content and neuropsychiatric disease. Dialogues Clin Neurosci. 2012;14(4):345–367. doi:10.31887/DCNS.2012.14.4/gbuzsaki

12. Başar E, Güntekin B. A review of brain oscillations in cognitive disorders and the role of neurotransmitters. Brain Res. 2008;1235:172–193. doi:10.1016/j.brainres.2008.06.103

13. Uhlhaas PJ, Singer W. Neural synchrony in brain disorders: relevance for cognitive dysfunctions and pathophysiology. Neuron. 2006;52(1):155–168. doi:10.1016/j.neuron.2006.09.020

14. Chen SX, Kim AN, Peters AJ, Komiyama T. Subtype-specific plasticity of inhibitory circuits in motor cortex during motor learning. Nat Neurosci. 2015;18(8):1109–1115. doi:10.1038/nn.4049

15. Kolar K, Dondorp D, Zwiggelaar JC, Høyer J, Chatzigeorgiou M. Mesmerize is a dynamically adaptable user-friendly analysis platform for 2D and 3D calcium imaging data. Nat Commun. 2021;12(1):6569. doi:10.1038/s41467-021-26550-y

16. Kleinfeld D, Giovannucci A, Friedrich J, et al. CaImAn an open source tool for scalable calcium imaging data analysis. Elife. Published online 2019. doi:10.7554/eLife.38173.001

17. Chukwu JE, Congdon EE, Sigurdsson EM, Kong XP. Structural characterization of monoclonal antibodies targeting C-terminal Ser404 region of phosphorylated tau protein. MAbs. 2019;11(3):477–488. doi:10.1080/19420862.2019.1574530

18. Zott B, Simon MM, Hong W, et al. A Vicious Cycle of b Amyloid-Dependent Neuronal Hyperactivation. http://science.sciencemag.org/

19. Busche MA, Grienberger C, Keskin AD, et al. Decreased amyloid-β and increased neuronal hyperactivity by immunotherapy in Alzheimer’s models. Nat Neurosci. 2015;18(12):1725–1727. doi:10.1038/nn.4163

20. Grienberger C, Rochefort NL, Adelsberger H, et al. Staged decline of neuronal function in vivo in an animal model of Alzheimer’s disease. Nat Commun. 2012;3. doi:10.1038/ncomms1783

21. Ko H, Hofer SB, Pichler B, Buchanan KA, Sjöström PJ, Mrsic-Flogel TD. Functional specificity of local synaptic connections in neocortical networks. Nature. 2011;473(7345):87–91. doi:10.1038/nature09880

22. Kuba H, Oichi Y, Ohmori H. Presynaptic activity regulates Na(+) channel distribution at the axon initial segment. Nature. 2010;465(7301):1075–1078. doi:10.1038/nature09087

23. Busche MA, Eichhoff G, Adelsberger H, et al. Clusters of hyperactive neurons near amyloid plaques in a mouse model of Alzheimer’s disease. Science. 2008;321(5896):1686–1689. doi:10.1126/science.1162844

24. Zott B, Simon MM, Hong W, et al. A vicious cycle of β amyloid–dependent neuronal hyperactivation. Science (1979). 2019;365(6453):559–565. doi:10.1126/science.aay0198

25. Menkes-Caspi N, Yamin HG, Kellner V, Spires-Jones TL, Cohen D, Stern EA. Pathological tau disrupts ongoing network activity. Neuron. 2015;85(5):959–966. doi:10.1016/j.neuron.2015.01.025

26. Green C, Sydow A, Vogel S, et al. Functional networks are impaired by elevated tau-protein but reversible in a regulatable Alzheimer’s disease mouse model. Mol Neurodegener. 2019;14(1). doi:10.1186/s13024-019-0316-6

27. Zhou L, McInnes J, Wierda K, et al. Tau association with synaptic vesicles causes presynaptic dysfunction. Nat Commun. 2017;8:15295. doi:10.1038/ncomms15295

28. McInnes J, Wierda K, Snellinx A, et al. Synaptogyrin-3 Mediates Presynaptic Dysfunction Induced by Tau. Neuron. 2018;97(4):823–835.e8. doi:10.1016/J.NEURON.2018.01.022

29. Decker JM, Krüger L, Sydow A, et al. Pro-aggregant Tau impairs mossy fiber plasticity due to structural changes and Ca(++) dysregulation. Acta Neuropathol Commun. 2015;3:23. doi:10.1186/s40478-015-0193-3

30. Polydoro M, Dzhala VI, Pooler AM, et al. Soluble pathological tau in the entorhinal cortex leads to presynaptic deficits in an early Alzheimer’s disease model. Acta Neuropathol. 2014;127(2):257–270. doi:10.1007/s00401-013-1215-5

31. Combs B, Christensen KR, Richards C, et al. Frontotemporal Lobar Dementia Mutant Tau Impairs Axonal Transport through a Protein Phosphatase 1γ-Dependent Mechanism. J Neurosci. 2021;41(45):9431–9451. doi:10.1523/JNEUROSCI.1914-20.2021

32. Alfaro-Ruiz R, Aguado C, Martín-Belmonte A, et al. Alteration in the Synaptic and Extrasynaptic Organization of AMPA Receptors in the Hippocampus of P301S Tau Transgenic Mice. Int J Mol Sci. 2022;23(21). doi:10.3390/ijms232113527

33. Prikas E, Paric E, Asih PR, et al. Tau target identification reveals NSF-dependent effects on AMPA receptor trafficking and memory formation. EMBO J. 2022;41(18):e10242. doi:10.15252/embj.2021110242

34. Miyamoto T, Stein L, Thomas R, et al. Phosphorylation of tau at Y18, but not tau-fyn binding, is required for tau to modulate NMDA receptor-dependent excitotoxicity in primary neuronal culture. Mol Neurodegener. 2017;12(1):41. doi:10.1186/s13024-017-0176-x

35. Suzuki M, Kimura T. Microtubule-associated tau contributes to intra-dendritic trafficking of AMPA receptors in multiple ways. Neurosci Lett. 2017;653:276–282. doi:10.1016/J.NEULET.2017.05.056

36. Hoover BR, Reed MN, Su J, et al. Tau mislocalization to dendritic spines mediates synaptic dysfunction independently of neurodegeneration. Neuron. 2010;68(6):1067–1081. doi:10.1016/j.neuron.2010.11.030

37. Decker JM, Krüger L, Sydow A, et al. The Tau/A152T mutation, a risk factor for frontotemporal-spectrum disorders, leads to NR2B receptor-mediated excitotoxicity. EMBO Rep. 2016;17(4):552–569. doi:10.15252/embr.201541439

38. Warmus BA, Sekar DR, McCutchen E, et al. Tau-Mediated NMDA Receptor Impairment Underlies Dysfunction of a Selectively Vulnerable Network in a Mouse Model of Frontotemporal Dementia. The Journal of Neuroscience. 2014;34(49):16482–16495. doi:10.1523/JNEUROSCI.3418-14.2014

39. Siddhartha MR, Anahi SG, Perla GP, et al. Phosphorylation of Tau protein correlates with changes in hippocampal theta oscillations and reduces hippocampal excitability in Alzheimer’s model. Journal of Biological Chemistry. 2018;293(22):8462–8472. doi:10.1074/jbc.RA117.001187

40. Rupert DD, Shea SD. Parvalbumin-Positive Interneurons Regulate Cortical Sensory Plasticity in Adulthood and Development Through Shared Mechanisms. Front Neural Circuits. 2022;16:886629. doi:10.3389/fncir.2022.886629

41. Wu J, Aton SJ, Booth V, Zochowski M. Heterogeneous mechanisms for synchronization of networks of resonant neurons under different E/I balance regimes. Frontiers in network physiology. 2022;2:975951. doi:10.3389/fnetp.2022.975951

